# GWAS-Informed data integration and non-coding CRISPRi screen illuminate genetic etiology of bone mineral density

**DOI:** 10.1101/2024.03.19.585778

**Authors:** Mitchell Conery, James A. Pippin, Yadav Wagley, Khanh Trang, Matthew C. Pahl, David A. Villani, Lacey J. Favazzo, Cheryl L. Ackert-Bicknell, Michael J. Zuscik, Eugene Katsevich, Andrew D. Wells, Babette S. Zemel, Benjamin F. Voight, Kurt D. Hankenson, Alessandra Chesi, Struan F.A. Grant

**Author notes:** Joint Authorship. These authors contributed equally to this work (Mitchell Conery, James A. Pippin, Yadav Wagley, Khanh Trang). These authors jointly supervised this work (Benjamin F. Voight, Kurt D. Hankenson, Alessandra Chesi, Struan F.A. Grant). Corresponding author, Struan F.A. Grant.

## Abstract

Over 1,100 independent signals have been identified with genome-wide association studies (GWAS) for bone mineral density (BMD), a key risk factor for mortality-increasing fragility fractures; however, the effector gene(s) for most remain unknown. Informed by a variant-to-gene mapping strategy implicating 89 non-coding elements predicted to regulate osteoblast gene expression at BMD GWAS loci, we executed a single-cell CRISPRi screen in human fetal osteoblasts (hFOBs). The BMD relevance of hFOBs was supported by heritability enrichment from stratified LD-score regression involving 98 cell types grouped into 15 tissues. 23 genes showed perturbation in the screen, with four (ARID5B, CC2D1B, EIF4G2, and NCOA3) exhibiting consistent effects upon siRNA knockdown on three measures of osteoblast maturation and mineralization. Lastly, additional heritability enrichments, genetic correlations, and multi-trait fine-mapping revealed unexpectedly that many BMD GWAS signals are pleiotropic and likely mediate their effects via non-bone tissues. Extending our CRISPRi screening approach to these tissues could play a key role in fully elucidating the etiology of BMD.

## INTRODUCTION

Low-trauma fragility fractures are a significant and common cause of increased mortality and morbidity in old age^1–3^. Low bone mineral density (BMD), a highly heritable^4–6^ and polygenic^7,8^ trait, is among the most important risk factors for such fractures^9^. This key trait has been the primary focus of genomic research into fracture etiology. Genome wide association studies (GWAS) have identified over 1,100 signals associated with BMD^8^, with each representing a possible therapeutic target to treat low BMD. Yet, there remains a fundamental obstacle in the conversion of BMD GWAS discoveries into new treatments, namely the identity of the underlying causal effector genes. Most GWAS loci (~90%)^10–12^ detect non-coding variant associations that likely confer their effects by altering the expression of nearby genes^13–15^, though which genes is often less than obvious.

Two interrelated issues have impeded broad identification of effector genes at non-coding BMD GWAS loci: 1) the development of a highly-parallelized screening technique capable of linking GWAS variants to their causal genes and 2) the need to determine the cellular and/or tissue context(s) relevant to each locus. Recently, CRISPRi screens targeted to non-coding elements have emerged as a powerful tool to solve the first of these issues. By pairing pooled CRISPRi perturbations of non-coding regulatory elements with single-cell RNA sequencing (scRNA-seq) readouts of gene expression, these screens can scale the identification of causal mediating effector genes to hundreds of loci in parallel without the need for artificial reporter constructs^16–30^. However, due to the technical requirement that any cell model used for a CRISPRi screen be easily transfected, these screens have thus far been applied in a limited number of disease contexts^17,21,22,26–30^, none obviously related to bone biology.

Regarding the second issue, while prior efforts have provided clear examples of BMD loci operating in specific cell types, most obviously in the osteoblast lineage responsible for bone deposition^31–34^, in general the full range of primary cell types that function in BMD pathophysiology remains uncharacterized. The only systematic genomic assessment of this issue was limited to cell types found in scRNA-seq of mouse bone^35^. Stratified linkage disequilibrium score regression (S-LDSC)^36^ offers an opportunity to determine BMD-relevant cell types throughout the whole body using human-derived measurements, specifically by finding cell types whose genomic regulatory regions are enriched for trait heritability.

In this work, we addressed both issues described above, providing a roadmap for elucidating effector genes across non-coding BMD GWAS loci through CRISPRi screening. Specifically, we leveraged S-LDSC to identify cell types and models relevant to BMD etiology, noting among the significant results a transfectable human osteoblast cell model, the human fetal osteoblast 1.19 cell (hFOB). We used this model to conduct a focused, pooled CRISPRi screen of 89 non-coding regions harboring putatively causal variants determined through linkage disequilibrium with GWAS sentinel SNPs and identified 23 perturbed genes. Using short interfering RNA (siRNA) knockdown, we then interrogated the roles of these genes in osteoblast differentiation and function, validating 15 with one or more significant osteoblast effects. Lastly, we corroborated additional S-LDSC heritability enrichments in metabolic and structural-tissue annotations by calculating cross-trait genetic correlations and conducting multi-trait fine-mapping. These analyses revealed that at both the genome-wide and locus-specific level, the genetic etiology of BMD relates to those of other cardiometabolic and anthropometric traits in ways that suggest a substantial proportion of BMD GWAS loci confer their effects in cell types beyond those of bone. This result has critical implications for future functional experiments designed to dissect the architecture of BMD and related disease endpoints.

## RESULTS

### S-LDSC identifies diverse cell types relevant to BMD

To identify potential model systems in which to execute a CRISPRi screen relevant to the etiology of BMD and fracture, we utilized genome-wide functional genomics to generate evidence supporting potential cell types relevant to BMD. Specifically, we applied stratified linkage disequilibrium score regression (S-LDSC)^36^ to partition the heritability of BMD within active and open chromatin regions annotated across a range of metabolic and structural cell types. As open and active chromatin generally reflects functional regulatory regions, cell types with enriched epigenetic annotations are more likely to play a role in BMD determination. For our analysis of public-domain and in-house-generated datasets, we employed the largest GWAS to date of BMD^8^ along with chromatin immunoprecipitation sequencing (ChIP-seq) peaks for activating histone marks (H3K27ac, H3K9ac, H3K4me1, and H3K4me3) and assay for transposase-accessible chromatin using sequencing (ATAC-seq) peaks to assess open chromatin. In total, we performed analysis on 210 genomic annotations across 98 primary cell types and models, which we grouped into 15 tissue categories (**Supplementary Table 1**).

After adjusting for multiple testing, we observed significant BMD heritability enrichment (Bonferroni adj. *P* < 0.05) across structural tissue annotations, namely those for osteoblasts, connective tissue, and skin (**Fig. 1**, **Supplementary Table 1**). Additionally, annotations for several metabolic tissues – adipose, cardiovascular, central nervous system, gastrointestinal, immune cells, liver, and skeletal muscle – also showed two or more significant enrichments. We then sought to determine whether there were any differences in the tissues relevant to BMD versus fracture by repeating the S-LDSC analysis on a GWAS of fracture incidence^8^ where we observed a broadly similar, but overall weaker pattern of heritability enrichment across tissue annotations (**Supplementary Fig. 1**).

**Figure 1.**
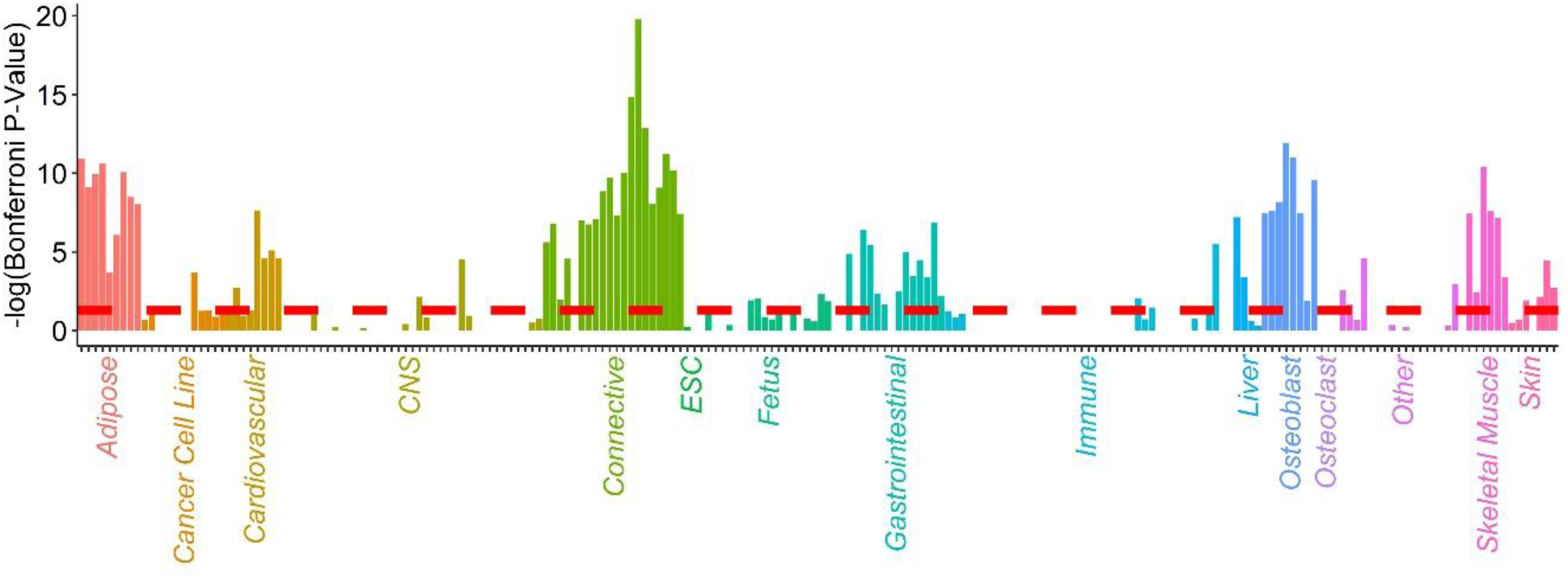
Bone mineral density partitioned heritability enrichments across 98 cell types. Each bar represents a particular genomic annotation (H3K27ac, H3K9ac, H3K4me1, H3K4me3, or open-chromatin) measured in a specific primary cell type or cell model. Negative log_10_ Bonferroni-adjusted p-values are plotted along the y-axis. The dashed line reflects a Bonferroni-adjusted significance cutoff of 0.05. Coloring reflects manually curated tissue categories for each cell type.

Unsurprisingly, given the known key role of osteoblasts in BMD determination, among the significant enrichments for BMD were several for primary osteoblasts as well as the hFOB and BMP2-stimulated human mesenchymal stem cell models (hMSC-osteoblasts) previously employed by ourselves^31,34,37,38^ and others^39–41^ to interrogate the genetic etiology of BMD in an osteoblast context. Specifically, we observed significant enrichment in primary-osteoblast H3K4me1 (adj. *P* = 1.15 × 10^−12^), H3K4me3 (adj. *P* = 1.01 × 10^−11^), and H3K27ac (adj. *P* = 6.81 × 10^−9^) ChIP-seq peaks in addition to those for H3K27ac in differentiated hFOBs (adj. *P* = 3.56 × 10^−8^). There was also enrichment in ATAC-seq peaks for differentiated hFOBs (adj. *P* = 0.013), 3-day differentiated pediatric hMSC-osteoblasts (adj. *P* = 3.23 × 10^−8^), 6-day differentiated pediatric hMSC-osteoblasts (adj. *P* = 2.38 × 10^−8^), and 3-day differentiated adult hMSC-osteoblasts (adj. *P* = 2.81 × 10^−10^).

However, we did not observe significant evidence for enrichment in the three annotations for osteoclast models included in our set (min. adj. *P* = 1) nor in the three annotations for monocytes (min. adj. *P* = 0.06), the direct precursors of osteoclasts. In fact, the only three annotations among the immune tissues to show enrichment were for hESC-derived CD56+ cultured mesoderm cells (adj. *P* = 3.27 × 10^−6^) and primary G-CSF-mobilized hematopoietic stem cells (adj. *P* = 0.009 and 0.033), both of which represent early progenitors capable of differentiating into many different lineages aside from the monocyte-osteoclast lineage.

### hFOB CRISPRi screen nominates effector genes at distal BMD GWAS loci

Given the enrichment of BMD heritability within the regulatory regions of primary osteoblasts and their corresponding cell models, we designed an osteoblast-focused screen leveraging the easily passaged and highly transfectable hFOB model. Our objective was to elucidate novel BMD effector genes for difficult-to-resolve GWAS signals mediated through distal regulatory effects rather than nonsynonymous coding variation or the disruption of gene promoters. Building on our prior experience with 3D genomic approaches to the elucidation of GWAS signals for other common complex traits^42–45^, we identified 88 such candidate signals that wholly resided in open chromatin outside of any active gene promoter but which showed physical interactions, as determined by chromatin conformation capture, with the open promoters of expressed genes in the hFOB and/or hMSC-osteoblast models (**Fig. 2A**, **Supplementary Fig. 2**). After merging two pairs of closely localized signals that fell within the effective repressive range of CRISPRi and including three additional signals that our group previously reported as associated with pediatric bone accrual^37^, our final tally of targets screened was 89.

**Figure 2.**
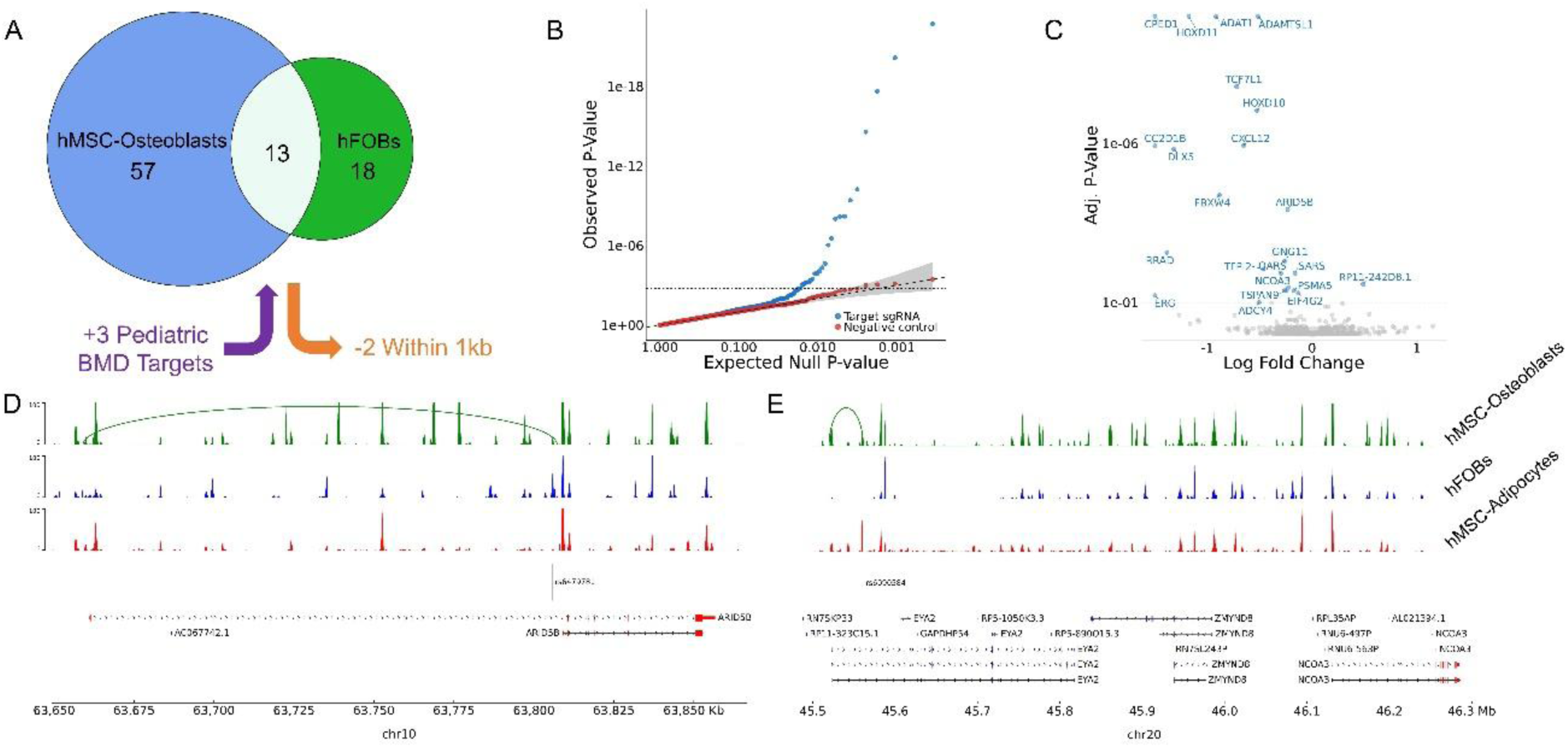
hFOB CRISPRi screen targets and perturbation results. (A) Breakdown of 89 screen targets by origin. (B) Quantile-quantile plot of targeting and non-targeting (negative-control) sgRNA tests. (C) Volcano plot of targeting sgRNA screen results. The 23 genes that exhibited significant perturbation at a target site are labeled. pyGenomeTracks plots of the (D) ARID5B and (E) EYA2 loci showing ATAC-seq read and Capture-C chromatin loops measured in hMSC-Osteoblasts, hFOBs, and hMSC-adipocytes. Targeted SNPs and genes are plotted below. All isoforms of genes perturbed in the CRISPRi screen are colored red.

We designed three synthetic guide RNAs (sgRNAs) for each of the 89 target sites and combined them into a pool with 27 scrambled non-targeting guides and two positive-control guides that targeted the transcription start sites of *RAB1A* and *SYVN1* (**Supplementary Table 2**). This pool was incorporated into lentiviral vectors and transfected into permissive hFOBs at a low multiplicity of infection (MOI) optimized to ensure that most viable cells received one sgRNA (see Methods). After differentiating and collecting the cells, we used scRNA-seq to determine both the gene expression profile of each cell as well the identities of any gRNAs they contained. Ultimately, we obtained 27,385 high-quality cells following filtering, each containing a single sgRNA (see Methods, **Supplementary Fig. 3-4**).

To verify the quality of the 27,385 cells, we investigated the expression of osteoblast marker genes within the retained population and observed robust expression of most though not all^46^ (**Supplementary Fig. 5**). To understand whether that result should have been expected, we compared pseudo-bulk expression of the 10,043 untargeted cells from our screen (2,340 that received only non-targeting guides and 7,703 that received no guide) with bulk RNA-seq results of differentiated and undifferentiated hFOBs and hMSC-Osteoblasts. Notably, we saw that the two sets of untargeted hFOBs from the screen had similar expression that was distinct from any of the bulk RNA-seq results (**Supplementary Fig. 6A**). Marker gene expression for the untargeted screen cells was comparable to that of the differentiated bulk hFOBs except for *ALPL* and *BGN* which had appreciably lower expression in the untargeted screen cells and *SPP1* which had much higher expression in the screen cells (**Supplementary Fig. 6B**). *SPP1* is a known inflammation marker^47,48^, so the expression differences could be a response to the lentiviral transfections or the presence of CRISPRi machinery in the otherwise untargeted cells. Overall, we observed no striking quality issues that would disqualify the transfected hFOBs from serving as a relevant model in an initial screen of BMD effector genes subject to orthogonal validation.

All but one sgRNA was successfully transfected into one or more of the 27,385 cells. The one failed transfection was due to the given sgRNA being substantially underrepresented in the lentiviral pool used for the screen (**Supplementary Table 2**). The remaining sgRNAs, yielded a median of 90 cells per guide with a minimum of 9 and a maximum of 354 (**Supplementary Fig. 7**). To test for significant perturbations in our screen, we used SCEPTRE^49,50^ to analyze the expression of all genes within 1MB of each target. SCEPTRE uses a permutation test to compare expression in cells that received a guide for each target site against those that received a non-targeting guide. SCEPTRE delivered well-calibrated results with minimal statistical inflation (**Fig. 2B**). Since our target selection method was agnostic to any observed or predicted effect directions, we did not know a priori whether the targets were active enhancers or repressors, and we allowed for both options in our analysis.

Beginning with the positive controls, we observed one of two that significantly repressed its target TSS (*RAB1A*). We lacked power to detect an effect for the other positive control, which targeted *SYVN1*, given only ~15% of cells expressed the gene. At the 89 target sites, we found 23 instances where perturbation of a target significantly impacted the expression of a gene within 1Mb; these instances spanned 23 distinct genes and 20 of the 89 distinct sites (adj. *P* < 0.10; **Fig. 2C**, **Supplementary Table 3**). All these perturbations were repressive except for one involving *RP11-242D8.1*, which is consistent with nearly all targets acting as enhancers. Eight of the perturbed genes, for example *ARID5B* (**Fig. 2D**), were predicted to be regulated by their target site based on the observed chromatin-conformation capture physical interactions. Conversely, the remaining 15 perturbations, involving genes like *NCOA3*, whose expression was affected by targeting rs6090584, an intronic variant within *EYA2* (**Fig. 2E**), were not reflected in the physical interaction dataset.

### Knockdown of perturbed genes reveals effects on osteoblast function

We next sought to corroborate the function of the perturbed candidate genes in osteoblasts and validate their status as BMD modulators key to osteoblast biology. To accomplish this, we first examined the effect the 20 successful cis perturbations in the hFOB screen had on the expression of osteoblast marker genes^46^. We reasoned that any trans-perturbations of the marker genes were most likely mediated via the observed 23 cis effects. Using SCEPTRE, we generated well-calibrated results (**Supplementary Fig. 8A**) that identified 10 significant trans-effects on osteoblast markers (adj. *P* < 0.10) for seven distinct target sites (**Supplementary Fig. 8B, Supplementary Table 4**). Nine of the ten significant trans-perturbations resulted in repression of the associated osteoblast marker, and a majority (63.1%) of all tested trans-perturbations showed directional repression. The target site that perturbed *SARS*/*PSMA5* was the only one to show significant upregulation of a marker gene, specifically *BGN*. This site was also notable for being one of three – in addition to the targets regulating *NCOA3* and *ADAT1* – for which we observed two significant trans-perturbations of osteoblast markers. Taken together these results suggest that at least a subset of the 23 candidate genes impact core elements of osteoblast biology.

To probe this issue further, we next used siRNA to directly knock down the candidate effector genes and evaluated their loss of function on osteoblast maturation and mineralization in the hFOB and hMSC-osteoblast models. As a measure of maturation, we assayed for alkaline phosphatase (ALP) staining in both cell models, and to assess mineralization we used Alizarin red S (ARS) assays in the hMSC-osteoblast model; hFOBs do not synthesize sufficient mineralized matrix for this second assay. Prior to beginning the assays, we manually reviewed the perturbed genes from the screen settling on a list of 21 to be tested in the assays that did not contain targeted exonic variants and were repressed rather than upregulated by CRISPRi (see Methods).

After knocking-down expression of the 21 genes in hFOBs, we observed evidence of decreased ALP activity for 11 of the 21 genes relative to scrambled siRNA controls (Benjamini-Hochberg adjusted *P* < 0.05; **Fig. 3A** top panel, **Supplementary Fig. 9**, **Supplementary Table 5**). When we completed the comparable experiment in hMSC-osteoblasts, we observed that knockdown of six of the 21 genes repressed ALP activity (**Fig. 3A** second panel, **Supplementary Fig. 10**, **Supplementary Table 6**). This decrease in the number of significant results is likely due to differences in the experimental model setting, but overall, we observed similar mean fold-changes between the two experiments. Taken together, five of the 21 genes (*ARID5B*, *CC2D1B*, *EIF4G2*, *FAM118A*, and *NCOA3*) revealed reduced ALP activity levels consistently across both cell models, while *CXCL12* was the only repressed gene to show effects in the hMSC-osteoblasts but not the hFOBs. Seeking to understand whether there were differences between the effect of the siRNA knockdowns on ALP protein activity and gene expression, we also used quantitative polymerase chain reaction (qPCR) to assess *ALPL* expression changes in hMSC-osteoblasts. Although most siRNAs showed directional repression of *ALPL* expression (**Supplementary Fig. 11**, **Supplementary Table 7**), none of the effects were significant (adj. *P* < 0.05). qPCR of other osteoblast markers similarly yielded no significant results.

**Figure 3.**
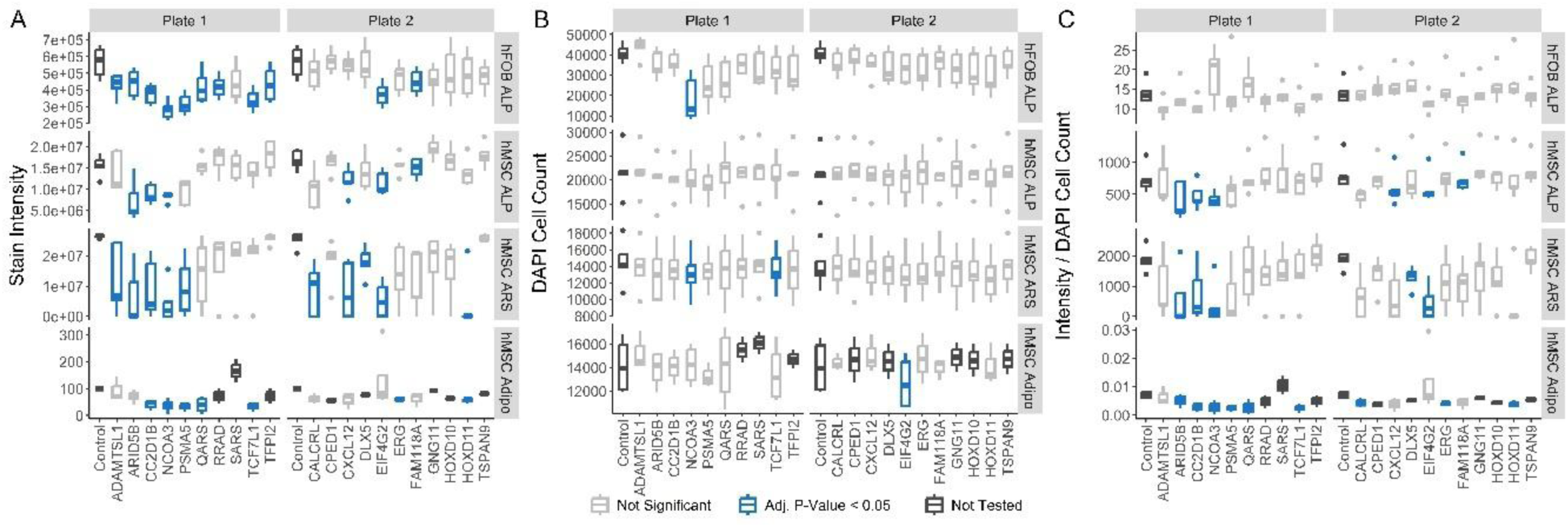
Assays of siRNA knockdown on osteoblast and adipocyte maturation and function. Results of Alkaline phosphatase (ALP) assay in hFOBs (top panels), ALP assay in hMSC-Osteoblasts (second from top), alizarin Red S (ARS) assay in hMSC-Osteoblasts (third from top), and adipogenesis assay in hMSCs (bottom) separated by (A) uncorrected stain intensity, (B) cell counts, and (C) per-cell stain intensity. In each plot siRNA targets are listed along the x-axis. hMSC-based assays were measured and normalized in the treated state (with BMP2 or adipogenic induction media). Any siRNAs resulting in significant decreases of assayed measurements are displayed in blue and siRNAs with < 3 replicates were untested and are marked in charcoal.

As for the ARS assay, the repression of ten genes led to decreases in the level of mineralization (**Fig. 3A** third panel, **Supplementary Fig. 12**, **Supplementary Table 8**). In total, the repression of four genes (*ARID5B*, *CC2D1B*, *EIF4G2* and *NCOA3*) showed consistent effects across the ALP and ARS assays, and 15 genes, showed effects on one or more of the osteoblastic phenotypes. Notable among the genes that showed no significant effects in the ALP and ARS assays were *HOXD10* and *CPED1*. These colorimetric assays taken together with results previously published by our team on the *CPED1*/*ING3* locus^31^, suggest that both *HOXD10* and *CPED1* represent instances where a gene without clear effects on BMD is co-regulated in osteoblasts by a GWAS-tagged regulatory element alongside a likely BMD-modulating gene, in these cases, *HOXD11* and *ING3* respectively.

These ALP and ARS assays are useful to identify genes with effects on osteoblast biology, but they cannot distinguish between genes that impact osteoblast viability or proliferation and those that impact cellular metabolism. To begin disentangling these issues, we re-stained each plate with DAPI to count the nuclei present in each well. DAPI staining was compatible with the ALP and ARS assays and no fluorescence interference was detected (**Supplementary Fig. 13**). From the DAPI staining, we directly analyzed the effects of each siRNA knockdown on cell count (**Fig. 3B**) and calculated the effects on the per-cell level of ALP activity and mineral secretion (**Fig. 3C**).

We observed relatively few instances of a gene knockdown resulting in a significant decrease in cell count. The notable exception was *NCOA3* which showed a drop in cell count with siRNA knockdown in both the hFOB ALP (**Supplementary Table 5**, adj. *P* = 0.03) and hMSC-Osteoblast ARS assays (**Supplementary Table 6**, adj. *P* = 0.03). *TCF7L1* knockdown also produced a slight cell-count decrease in the ARS assay (**Supplementary Table 8**, adj. *P* = 0.046). In contrast, we observed more significant effects of siRNA knockdown on the cell-count normalized assay results, though only in the hMSC assays. Among the significant hits, knockdown of *ARID5B*, *CC2D1B*, *EIF4G2* and *NCOA3* all produced decreases in hMSC-osteoblast ALP activity (**Supplementary Table 6**) and mineral secretion (**Supplementary Table 8**, adj. *P* < 0.05). Taken together, the cell-count and per-cell assay results provide nuance to several observations from the overall osteoblast assays. For example, they indicate that the four genes with consistent effects in the main assay results (*ARID5B*, *CC2D1B*, *EIF4G2*, and *NCOA3*) likely all affect per-osteoblast levels of bone deposition while simultaneously suggesting that *NCOA3* may play an additional role in maintaining osteoblast viability or proliferation. However, in many cases, the main assay results cannot be clearly resolved into specific impacts on cell-count and per-cell activity.

To support the results of these osteoblast experiments, we attempted to confirm the effectiveness of the siRNA knockdowns in the hMSC model, however, it appears we lacked power to detect significant knockdown for many siRNAs using the same donor cell lines as in the assays. At 4 days post differentiation, we detected significant repression (adj. *P* < 0.05) only for *CPED1*, *NCOA3*, *QARS* and *TSPAN9* in the hMSC-osteoblasts, but observed similar magnitude expression decreases for nearly all other genes (**Supplementary Fig. 14A**, **Supplementary Table 9**). *ARID5B* and *CC2D1B* were notable exceptions with the smallest median decreases in expression (12% and 26% respectively) among 17 tested genes. We also failed to evaluate four genes for knockdown validation (*FAM118A*, *HOXD10*, *HOXD11*, and *SARS*) due to inconsistent amplification of PCR primers across hMSC donors.

In experiments in which assays are replicated on multiple plates, it is often standard to normalize each well to the control well from the same plate to remove confounding inter-plate variability unrelated to the perturbations being examined. We recalculated all our assay results from the raw data (**Supplementary Tables 10-11**) using such plate-normalization (**Supplementary Fig. 15**, **Supplementary Tables 12-14**). However, having observed relatively few differences, particularly for the four genes with consistent osteoblast effects (*ARID5B*, *CC2D1B*, *EIF4G2*, and *NCOA3*), which remained unaffected in the main analysis, we report the more conservative results obtained without this added normalization step.

### Certain BMD candidate effector genes play an adipogenic role

To elucidate how knockdown of the genes with observed effects disrupted osteoblast function, we also assessed whether siRNA knockdown would restrict the ability of hMSCs to differentiate along the adipocyte trajectory, an alternative lineage to the osteoblast/osteocyte path. We hypothesized that if the suppression of any genes also disrupted adipogenesis, it would indicate that those genes are involved in upstream pathways that regulate the switch between hMSC proliferation and differentiation. Given the long timeline of adipogenic differentiation and availability of stocks for some donor lines, we ran these assays in two steps, first testing effects in two hMSC lines followed by additional replicates and significance testing for genes with appreciable reductions of intracellular lipid droplets in the initial lines and significant results in the hMSC-osteoblast assays. Combining replicates across the two steps, seven siRNAs significantly impaired the adipogenic potential of the hMSCs (adj. *P*-value < 0.05; **Fig. 3A** bottom panel, **Supplementary Fig. 16**, **Supplementary Table 15**). The seven genes with significant adipogenic effects remained unchanged by plate-normalization (**Supplementary Fig. 15**, **Supplementary Table 16**) and all remained significant when examining per-cell adipogenic effects (**Fig. 3C** bottom panel, **Supplementary Table 15**).

Focusing on the four genes that had consistent effects in the colorimetric osteoblast assays, we observed that the results of their knockdown aligned with the genetic architecture observed at their loci in osteoblast and adipocyte cell models. For example, *NCOA3* and *CC2D1B* repression impaired adipogenesis, and the target site for each intersected an ATAC-seq peak found in both osteoblast and adipocyte cell models (**Fig. 2E** and **Supplementary Fig. 17**). In contrast, the *ARID5B*-linked target resides within osteoblast-specific open chromatin (**Fig. 2D**), and the *EIF4G2*-linked target was observed to have an interaction with the *EIF4G2* promoter only in the hMSC-osteoblast model (**Supplementary Fig. 18**). However, the lack of an observed interaction in adipocytes may not be a sufficient explanation for *EIF4G2*’s lack of effect on adipogenesis as a similar hMSC-osteoblast-specific interaction was observed at the *TCF7L1* locus (**Supplementary Fig. 19**), though adipogenic effects were detected for *TCF7L1* siRNA knockdown.

As before, we also attempted to validate the siRNA knockdown using the same number of donors selected in the two-step assay process for each gene. We again observed four significantly repressed genes (*CALCRL*, *NCOA3*, *QARS*, and *TCF7L1*) at 12 days post-differentiation (**Supplementary Fig. 14B**, **Supplementary Table 17**). Unfortunately, there were two genes, *CC2D1B* and *RRAD*, that showed no directional repression in differentiated adipocytes, and *RRAD* was similarly unrepressed in undifferentiated hMSCs. However, most genes showed directional repression in both the differentiated and undifferentiated hMSCs with similar median percentage changes in expression to the significantly repressed genes.

### Pleiotropic genetic architecture for BMD supports metabolic tissue enrichments

As mentioned above, our primary objective was to validate BMD genes at non-coding GWAS loci using a CRISPRi screening system. To do so, we leveraged the most obvious cell type with BMD heritability enrichment, i.e. the osteoblast lineage. However, given we had successfully implicated effector genes using our strategy in this initial cellular setting, we returned to consider the ramifications of our presented S-LDSC analyses described above, where we observed potential roles for several metabolic and structural cell types in BMD determination. If valid, these observations suggest that to fully characterize the entire genetic architecture at BMD GWAS loci, non-coding CRISPRi screens and other orthogonal functional approaches would need to be applied across a range of cell models, and crucially beyond those traditionally considered directly relevant to bone biology. We reasoned that if these additional tissues are critical to BMD pathophysiology, then BMD should share genetic etiology with phenotypes known to be related to them. To this end, we systematically investigated how the etiology of BMD relates across 37 other anthropometric and cardiometabolic traits using publicly-available GWAS conducted in the UK Biobank^8,51^ (**Supplementary Table 18**).

We first investigated the cross-trait relationships of BMD at the genome-wide level via LDSC-based genetic correlations. In total, we detected a significant correlation (Benjamini-Hochberg adj. *P* < 0.05) between BMD and eleven additional traits (**Fig. 4** and **Supplementary Table 19**). As expected, we observed a strong positive correlation of the trait with itself and an inverse correlation with the incidence of bone fracture. We also validated a recent result showing an inverse relationship with sex-hormone binding globulin^52^. Many of the remaining significant correlations were for traits related to body composition and fat including body mass index (r_g_ = 0.07), body fat percentage (r_g_ = 0.05), trunk fat percentage (r_g_ = 0.04), whole-body impedance (r_g_ = −0.06), and whole-body fat mass (r_g_ = 0.05). These correlations align well with the observed heritability enrichments in adipose, but it should be noted that they are all derived from similar anthropometric measurements and are all correlated with one another (**Supplementary Fig. 20** and **Supplementary Table 20**).

**Figure 4.**
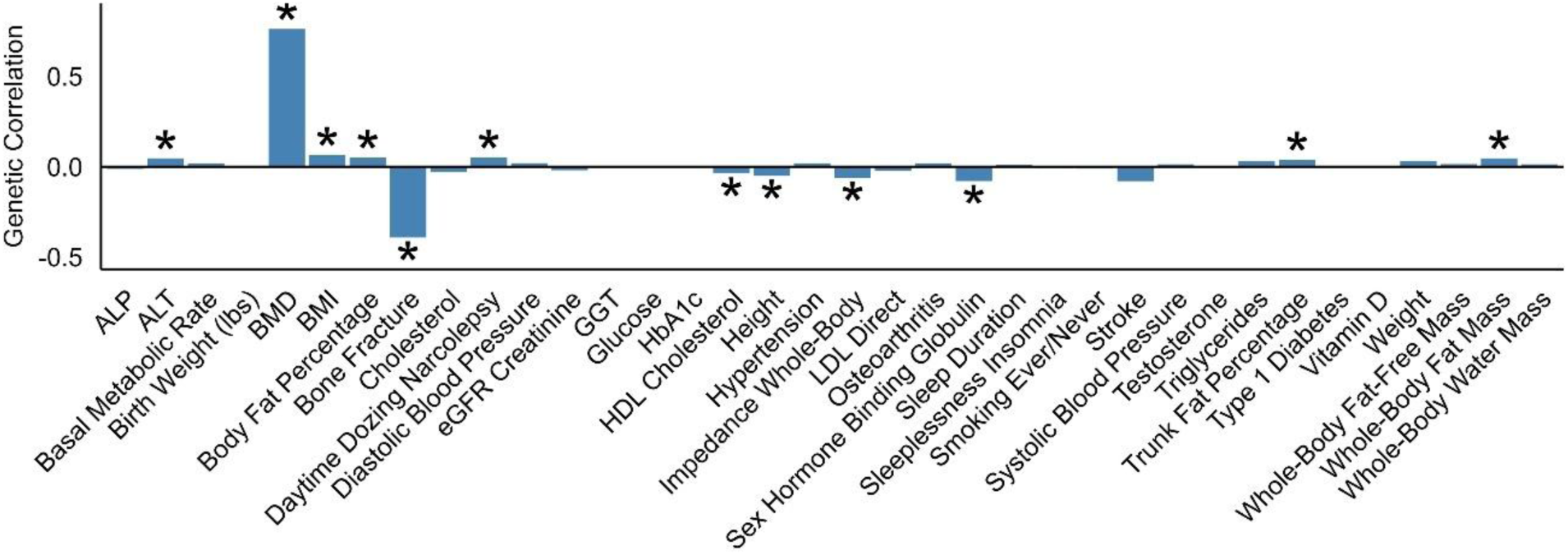
Genetic correlations between BMD and 38 traits. Genetic correlations are displayed the vertical axis and traits along the horizontal. The 38 traits include the correlation of BMD with itself. Significant correlations (Benjamini-Hochberg Adj. *P*-Value < 0.05) are labeled with an asterisk.

We further investigated the shared cross-trait etiology of BMD at the locus-specific level using CAFEH (colocalization and fine-mapping in the presence of allelic heterogeneity)^53^ and an approximate in-sample linkage disequilibrium (LD) reference panel to conservatively fine-map and colocalize 433 independent BMD signals, which we presumed captured at least one BMD-modulating variant each (see Methods, **Supplementary Tables 21-23**). 123 of the BMD signals were shared with one or more other traits, with 22 of the 37 input traits mapped to one or more signals (**Supplementary Fig. 21**). The greatest number of signals were shared with height (42 signals); three body-composition and weight-related traits: whole-body water mass (27 signals), whole-body fat-free mass (26 signals), and basal metabolic rate (25 signals); and serum ALP (22 signals).

Using hierarchical clustering to group the signals by the traits for which each was mapped, we found a cluster of signals mapped to body-composition traits that suggested a shared genetic etiology (**Fig. 5A**). Included among this cluster was the single most pleiotropic signal which consisted of a single variant at the *CCND2* locus, rs76895963 (MAF = 2.1%), and which mapped to 13 traits including BMD (**Supplementary Fig. 22**). There were also distinct groups of signals linked to (i) height independent of body-composition, (ii) serum ALP, (iii) blood pressure, and (iv) BMD only. Visualization with uniform manifold approximation projections (UMAP)^54^ yielded a similar clustering of signals, with observed clusters related to body composition, body-composition-independent height, ALP, blood pressure, and BMD only (**Supplementary Fig. 23**). Seeking to understand whether these clusters could reflect underlying pathways, we examined the effect directions of the signals across the traits (**Supplementary Table 24**). In most cases, the signals identified for any trait were split between those that had positively correlated effects on the trait and BMD and those with negatively correlated effects (**Fig. 5B**). A notable exception was bone fracture incidence for which all six mapped signals had effects that correlated negatively with BMD. When re-clustering the signals accounting for effect directions, we observed that the major clusters all split, indicating there could be multiple pathways relating groups of traits, some with counteracting effects (**Supplementary Fig. 24**). In summary, these results taken together with the S-LDSC heritability enrichments, support a complex model of BMD genetic architecture that is both pleiotropic across a large subset of GWAS loci and mediated by multiple distinct pathways and cell types.

**Figure 5.**
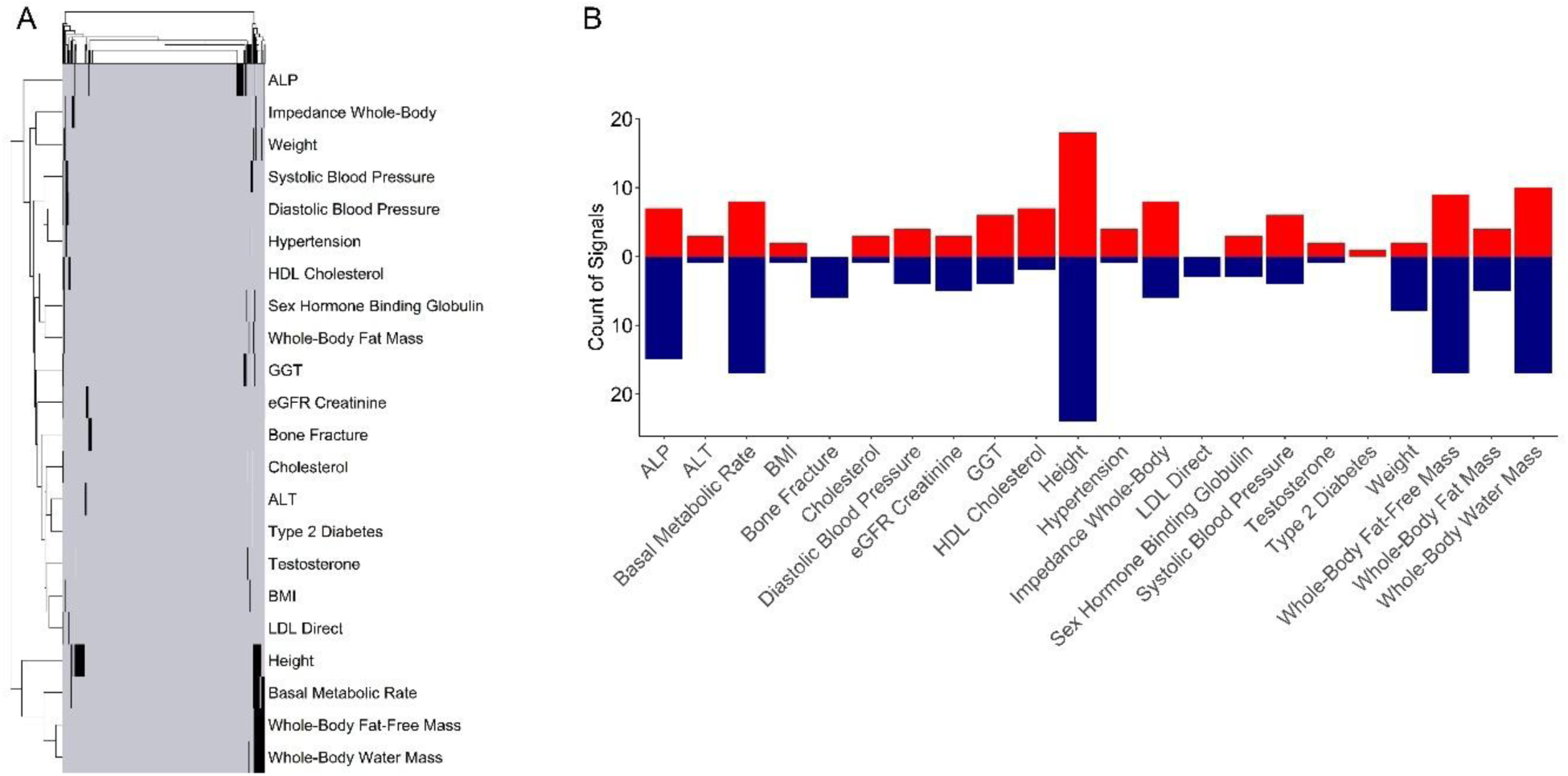
Cross-trait signal sharing and effect directions of 433 BMD signals across 22 traits. (A) Heatmap and dendrogram of trait mappings. Traits are plotted down the y-axis with individual signals plotted along the x-axis. Black coloring indicates when a signal modulates the corresponding trait. Signals and traits are clustered hierarchically using the complete-linkage method with Euclidean distances. (B) Breakdown of signals mapped per trait split into signals with positively correlated effects on BMD and given trait (red) and signals with negatively correlated effects (blue).

## DISCUSSION

In this study, we demonstrate the power of non-coding CRISPRi screens to elucidate novel regulatory elements at BMD GWAS loci implicating 23 putative BMD-modulating genes, 15 of which we later validated *in vitro* to affect one or more measures of osteoblast maturation or mineralization in hFOBs and/or hMSC-osteoblasts. Notably, we found down-regulation of four genes (*ARID5B*, *CC2D1B*, *EIF4G2*, and *NCOA3*) exhibited directionally-consistent repressive effects in all three osteoblast-focused assays. Additionally, we showed that knockdown of two of them, *CC2D1B* and *NCOA3*, also impaired differentiation of hMSCs into terminal adipocytes which aligns with genomic evidence that these genes may impact BMD determination by acting at an earlier time point in hMSC differentiation. Lastly, we observed S-LDSC heritability enrichments, genetic correlations, and multi-trait fine-mapped signals as evidence that a plethora of metabolic and structural cell types and widespread pleiotropic inheritance are critical to BMD etiology. This underlines the future challenge of applying CRISPRi screens and other experimental techniques to the complete resolution of causal effector gene identities at BMD GWAS loci.

Regarding our design, we were surprised to observe that only 8 of the 20 significantly-perturbed target sites (42.8%) were observed to regulate one of the predicted genes with whose promoters they interacted in the hFOB and hMSC-osteoblast Capture-C datasets. While this proportion was higher than the 23.9% (32/134) of significantly perturbed targets in a recent screen of K562 cells that were reflected by H3K27ac HiChIP contacts^21^, it failed to include three genes – *KARS*, *TEAD4*, and *PRPF38A* – which we previously observed to have osteoblastic effects and are regulated by enhancers at the included pediatric bone accrual loci^37^. Although this may reflect temporal activities at given loci, this discordance most likely reflects differences in sensitivities between Capture-C and CRISPRi screens and highlights the value of embracing a “confluence of evidence” to nominate putative causal genes. Fortunately, these differences in sensitivity provided the opportunity to nominate *CC2D1B* as a second putative BMD gene at the locus harboring *PRPF38A*. These two genes provide an interesting example of a GWAS signal tagging multiple co-regulated genes with independent effects on osteoblast function, and likely BMD. In contrast, between this study and our prior work^31^, we have also observed two loci with co-regulated genes where only one gene has observed osteoblastic effects, namely, *HOXD10*/*HOXD11* and *ING3*/*CPED1*. That we observed evidence for both patterns of co-regulation emphasizes the need to validate putative genes derived from CRISPRi screens or any other nomination schema in orthogonal assays.

The four BMD candidate genes best supported by our results have various levels of prior evidence in the published literature. The strongest support in both our work and the literature is for *NCOA3*, which has previously been shown to impact BMD through a polyglutamine repeat expansion located at the protein’s carboxy-terminal^55,56^. *NCOA3* knockout has also been linked to hearing loss and altered bone and cartilage formation in zebrafish^57^ as well as effects on chondrocyte formation and osteoarthritis through rs6094710, a non-coding variant 35kb upstream of *NCOA3*’s transcription start site. However, our work appears to be the first implicating an effect on *NCOA3* expression for rs6090584, which is in low LD with rs6094710 (R^2^ = 0.002) and located more than 500kb upstream of *NCOA3*. *ARID5B* and *EIF4G2* have more indirect support in the literature as putative BMD effector genes. Knocking out *ARID5B* leads to mouse skeletal abnormalities^58–60^. Additionally, during chondrogenesis, a process closely related to bone formation, *ARID5B* and its homolog, *ARID5A*, act as physically-interacting coactivators of *SOX9* recruiting histone demethylases and acetylases respectively^61,62^. *EIF4G2* and the multiple miRNAs that regulate it^63–66^ have been shown to impact chondrogenesis^65^, the repair of nucleus pulposus cells^64^, the activity of osteosarcoma cells^66^, and most recently, osteoblastogenesis^63^. In contrast to the other three genes, *CC2D1B* has seemingly never been implicated in bone biology. The most closely related work has shown upregulation of *CC2D1B* in periodontal ligament cells subjected to orthodontic force^67^, but most research focused on the gene has probed its numerous roles across cell types in the endosomal sorting complex required for transport (ESCRT) machinery^68–71^.

Two of the most surprising results from this study were the observed metabolic and structural-tissue heritability enrichments; and the lack of enrichment in the tested osteoclast annotations. Of course, we acknowledge that not all cell types with enriched epigenetic annotations are necessarily causal for BMD. Spurious enrichments can result from correlations of epigenetic features with true causal cell types via shared regulatory pathways or linkage disequilibrium with causal variants active in distinct cell types. We are aware of a few methods that leverage expression quantitative trait loci (eQTL) to fine-map causal cell-types^72–74^ but were cautious to employ them as they are limited by the systematic differences between the discoverability of GWAS and eQTL signals^75^ and the availability of bone-cell eQTLs, which is restricted to a dataset of primary osteoblasts from surgical explants^76^ and a dataset of RANKL-stimulated osteoclast-like cells^77^. Instead, we chose to interpret our S-LDSC results principally at the lower-resolution of tissues, which should be less susceptible to spurious correlations between closely related cell types.

Moreover, we have identified genetic correlations and multi-trait fine-mapped signals that support the heritability enrichments. To illustrate, the cardiovascular tissue enrichments could plausibly be linked to the small cluster of blood pressure signals, and the cluster of ALP signals coincides with the observed heritability enrichments in liver, the other major source of serum ALP besides bone^78^. More obviously, adipose tissue enrichments align well with positive genetic correlations between BMD and body-fat percentage, BMI, and whole-body fat mass. Similarly, the cluster of signals fine-mapped predominantly to basal metabolic rate and whole-body fat-free mass support the observed enrichments in skeletal muscle tissue. Much research has focused on the relationships between body composition and BMD. Briefly, muscle mass, strength, and bone density have generally been found to correlate positively across groups^4,79–85^ likely sharing a causal relationship^86–88^, while the association between adiposity and BMD is more complex with some evidence of positive correlations between adiposity and the BMD of weight-bearing bones^83,85,86,89,90^ that may attenuate or even reverse after accounting for adiposity type^84,91–93^, at extreme ranges of adiposity^85,93,94^, or in certain age and sex-based sub-populations^83,86,92,95^. The complexity of these prior results coincides well with our observation of effect direction heterogeneity across signals fine-mapped for both body-composition traits and BMD. Such heterogeneity could also explain the small magnitudes observed for genetic correlations with body-composition traits though more work would need to be done to confirm this hypothesis and fully elucidate the clearly multi-faceted relationship between body composition, size, and BMD. Given the insight available at this point, it is only obvious that these traits are somehow related, and therefore it is logical that at least a subset of BMD GWAS loci mediate these relationships in cell types related to body composition.

In contrast, the lack of BMD heritability enrichment in osteoclast annotations and those of their predecessors, monocytes, seems counterintuitive given osteoclasts’ critical role in remodeling bone and since disrupted osteoclast activity has been proposed to mediate several specific BMD GWAS loci^77,96–98^. However, as we calculated heritability enrichments on top of the baseline S-LDSC model which accounts for conservation and genomic regulation broadly relevant across cell types, a lack of enrichment in osteoclast annotations does not prohibit isolated BMD GWAS loci from mediating their effects in osteoclasts, it merely suggests that in general, osteoclasts are not as critical to the etiology of BMD as the other enriched cell types. In fact, there may be some orthogonal evidence for this conclusion. A recent study leveraging scRNA-seq of mouse bone and a variant of S-LDSC^99^ found a similar lack of heritability enrichment in osteoclasts^35^ and another unrelated study observed that a far larger proportion of BMD-associated SNPs were eQTLs in adipose and skeletal muscle than in osteoclasts^100^. Of course, other explanations for the lack of enrichment are also possible. It may be that osteoclasts are more critical to BMD in its extreme ranges, such as in osteoporosis patients, but that osteoclast activity is less relevant to BMD variation in healthy individuals, such as those that comprise most of the UK Biobank cohort. Similarly, it could be possible that the RANKL-stimulated osteoclast cell model used for the osteoclast annotations does not match the primary cell closely enough to detect enrichment, perhaps due to an insufficient duration of differentiation. However, detracting from this latter hypothesis, all three osteoclast regulatory annotations were found to strongly overlap the promoters of osteoclast marker genes^101^ (see Methods).

There are several key limitations of our work in both experimental and bioinformatic respects. Firstly, on the experimental side, the CRISPRi screen was conducted in an osteoblast cell model, which may not fully reflect primary cell activity. In fact, it had low expression of a few key osteoblast marker genes, possibly owing to an inflammatory response to the CRISPRi transfection. The hFOB and hMSC models used for the assays also differ from one another, and in instances where they provide conflicting results, as in the case of the cell-count normalized ALP assays, it is not clear which, if either, set of results is closer to primary osteoblast activity. Regarding the unnormalized results, technical factors may have also affected the cell counts independent of the siRNA knockdowns. Additionally, as we focused on osteoblast biology, it is possible that some of the targeted GWAS signals and nominated genes may have additional BMD-relevant effects in other cell types and tissues. Targeted variants with significant perturbations also cannot be assumed to be causal without validation due to the potential for CRISPRi-induced heterochromatin to span linked, true causal variants or even inhibit unrelated regulatory mechanisms. For example, the targeted variant at the *CC2D1B* locus, rs34455069, lies 430bp upstream of the *CC2D1B* promoter within the first intron, and the CRISPRi heterochromatin may have expanded into the promoter region blocking *CC2D1B* expression independent of rs34455069. This problem may have also occurred at the *HOXD10*/*HOXD11* locus where the targeted variant is 350bp upstream of the promoter. Both loci also contain examples of candidate effector genes for which we were unable to demonstrate significant siRNA repression. Though we observed mostly consistent directional knockdown of the targeted genes, our ability to detect significant siRNA knockdowns was impaired by weak and noisy effects, unreliable primers, and our supply of suitable hMSC lines. In a few instances, we even saw no directional repression by the siRNAs. These mixed qPCR results weaken the link between the candidate effector genes and BMD and leaves open the possibility that off-target knockdowns are responsible for some of the assay results. Our decision to purchase pooled siRNAs may have reduced mis-targeting as each siRNA in a pool is likely to have a unique set of spurious off-targets, however without being able to separate the siRNAs, we were unable to test and validate this hypothesis. Moreover, only having assessed siRNA knockdown at a single time point during the early processes of osteoblast and adipocyte differentiation, we cannot draw conclusions about the effectiveness of siRNA knockdown at later points of differentiation nor understand how gene repression at distinct points in the trajectory impact terminal phenotypes such as ARS staining and lipid droplet formation.

On the bioinformatic side, the S-LDSC enrichments, genetic correlations, and multi-trait fine-mappings depend on the power of their underlying GWAS and are affected by mismeasurement or heterogeneity of the phenotypes. Such phenotypic heterogeneity may explain the fewer enrichments observed in fracture incidence relative to BMD. The S-LDSC results are also constrained by the inconsistent availability of ATAC-seq and ChIP-seq datasets across cell types which could biasedly cause heavily profiled cell types to have multiple enrichments for biologically correlated ATAC-seq and ChIP-seq annotations and less profiled tissues to have fewer enrichments independent of BMD relevance. The computational results may also not be fully transferrable outside of individuals genetically similar to European reference populations^102^. Lastly, seeking to prioritize precision over recall, we tolerated a high-false negative rate in the CAFEH fine-mapping to ensure that the signals we identified and the traits to which we mapped them were well-supported. For this reason, the 433 signals we report are far fewer than the 1,103 conditionally-independent signals mapped by Morris *et al*. using the same GWAS^8^.

In summary, we have identified 23 putative causal genes of which we were able to demonstrate 15 had at least one osteoblast effect and provided strongest evidence for four: *ARID5B*, *CC2D1B*, *EIF4G2*, and *NCOA3*. This work demonstrated the power of non-coding CRISPRi screens in relevant cell models to elucidate unknown biology and causal genes at BMD GWAS loci. We also characterized the tissues relevant to BMD etiology and corroborated them via genetic correlations and multi-trait fine-mapped signals. Jointly, these results provide a roadmap for how this powerful experimental technique may be applied to the challenging task of resolving effector gene identities at all BMD GWAS loci.

## METHODS

### Acquisition and processing of hFOBs for ATAC-seq and Capture-C

hFOBs were purchased from ATCC and maintained in a permissive state at 33.5°C in a 1:1 mixture of Ham’s F12 Medium and Dulbecco’s Modified Eagle’s Medium with 2.5 mM L-glutamine (without phenol red), 10% Fetal Bovine Serum (FBS), and 0.3 mg/ml G418 sulfate solution. All experiments were performed on cells lower than passage 8 and confirmed to be mycoplasma negative. Cells were differentiated by increasing culture temperature to 39.5°C and were harvested for ATAC-seq and Capture-C five days post-differentiation. Matched undifferentiated control cells were also collected at the same time. Three biological replicates of the undifferentiated and differentiated hFOBs were collected for Capture-C with a fourth replicate of the differentiated hFOBs collected for ATAC-seq.

### Differentiation of primary human mononuclear cells into osteoclasts for ATAC-seq

Human bone marrow mononuclear cells purchased from Lonza were utilized to generate and characterize human osteoclasts with minor modifications as described by Cody *et al*.^103^ and Susa *et al*.^104^ Cells were cultured in alpha-MEM containing 10% FBS supplemented with 33 ng/ml recombinant M-CSF for 2 days before using for differentiation. For differentiation, 2 × 10^5^ cells were seeded onto a well of a 24 well plate and cultured in differentiation medium containing 33 ng/ml M-CSF, 66 ng/ml human RANKL and 1 ng/ml TGF-beta1. Medium containing supplements were re-fed every 3-4 days for a total of 12 days after which the cells were evaluated for morphological changes and stained for tartrate-resistant acid phosphatase (TRAP) with a commercially available leukocyte acid phosphatase kit (SIGMA, Cat. 387-A). Three replicates of differentiated cells were processed to prepare samples for ATAC-seq at 0, 4, 8, and 12 days.

### Acquisition of pediatric hMSCs and differentiation to osteoblasts for ATAC-seq

hMSCs were obtained from the surgical waste of six pediatric patients undergoing ACL reconstruction surgery at the MOTT Children’s Hospital, University of Michigan. Samples were processed following the protocol we published previously for adult hMSCs^31^. Briefly, the bone reamings were digested with collagenase for 3 hours and plated on a 10 cm dish. Cell colonies were lifted with Trypsin-EDTA and cell lines were established. Established lines were characterized by expression of MSC markers, and additionally tested for adipocyte, osteoblast, and chondrocyte differentiation. Validated cells were used for ATAC library generation. Cells were collected at 3 days and 6 days post BMP2-stimulated differentiation and at 3 days post mock stimulation.

### hFOB, pediatric osteoblast, and osteoclast ATAC-seq library generation

Fresh hFOBs, pediatric osteoblasts, and osteoclasts were harvested via Trypsin or TrypLE, followed by a series of DPBS wash steps. 50,000 cells from each sample were pelleted at 550 × g for 5 minutes at 4 °C. The cell pellet was then resuspended in 50 μl cold lysis buffer (10 mM Tris-HCl, pH 7.4, 10 mM NaCl, 3 mM MgCl2, 0.1% IGEPAL CA-630) and centrifuged immediately at 550 × g for 10 minutes at 4 °C. The nuclei were resuspended in transposition reaction mix (2x TD Buffer (Illumina Cat #FC-121–1030, Nextera), 2.5 µl Tn5 Transposase (Illumina, 20034197 Cat #FC-121–1030, Nextera) and Nuclease Free H2O) on ice and then incubated for 45 minutes at 37°C. The transposed DNA was then purified using the MinElute Kit (Qiagen), eluted with 10.5 μl elution buffer (EB), frozen and sent to the Center for Spatial and Functional Genomics at CHOP. The transposed DNA was PCR amplified and indexed using the Illumina Nextera Kit (Illumina) and NEBNext High-Fidelity 2x PCR Master Mix (NEB) for 12 cycles to generate each library. The PCR reaction was subsequently purified using AMPureXP beads (Agencourt) and libraries were paired-end sequenced on the Illumina NovaSeq 6000 platform.

### Human articular chondrocyte isolation

Human knee articular cartilage was provided by AlloSource (Centennial, CO) from donors deemed eligible for tissue donation for research purposes. Donor eligibility was determined in accordance with American Association of Tissue Banks (AATB) and Food and Drug Administration (FDA) regulations. Tissue fragments that included subchondral bone and cartilage from both the tibial plateau and femoral condyles were surgically removed from donors (N=3) and immediately placed into pre-chilled (wet ice) wash medium composed of DMEM/F12 medium (Cytiva, #SH30023.01) containing amphotericin (1 ng/ml, Sigma, #50-175-7519), gentamycin (0.05 µg/ml, Gibco, #15750-060), and Pen/Strep (1% v/v, VWR, # K952-100ML) for transport to the laboratory. Articular cartilage was removed from the subchondral bone using a scalpel, minced into 1 mm × 1 mm fragments in a Petrie dish, rinsed three times with phosphate-buffered saline, and placed into a 50 ml conical tube containing 30 ml wash medium supplemented with fetal bovine serum (FBS, 25% v/v, Gibco, #10438-026). Chondrocytes were isolated from the cartilage using a modified version of an established method^105^. Specifically, after 60 minutes of gentle shaking at 37°C, medium was aspirated and replaced with 30 ml digestion medium composed of DMEM/F12, 25% FBS, 0.05 µg/ml gentamycin, 1% v/v Pen/Strep, and ascorbic acid (100 µg/ml, Sigma, #A4544-25G), and supplemented with pronase (0.3 µg/ml, Roche, #10165921001). Cartilage fragments were digested in this medium for 90 minutes at 37°C with gentle shaking, followed by centrifugation at 300 × g at 4°C for 7 minutes to pellet tissues and cells, followed by resuspension in 30 ml digestion medium supplemented with collagenase II (1.2 mg/ml, Worthington, #LS004177). Cartilage was digested for 18 hours at 37°C with gentle shaking and filtered through a 70 µm strainer, with a single rinse of the tube with wash medium to recover all remaining cells and tissue, which was again strained. Strained materials were centrifuged to a pellet at 300 × g for 7 minutes at 4°C, resuspended with fresh wash medium, spun once more at 300 × g for 7 minutes at 4°C to form a pellet, and then finally resuspended in DMEM/F12 containing 20% FBS, 0.05 µg/ml gentamycin, 1% v/v Pen/Strep, and 100 µg/ml ascorbic acid. Isolated suspensions of chondrocytes were counted using a hemocytometer.

### Human chondrocyte nucleic acid preparation for ATAC-seq

Immediately following human articular chondrocyte isolation, nucleic acids were extracted for use in an ATAC-seq experiment. From each human donor (N=3), suspensions of 75,000 cells were centrifuged at 550 × g for 5 min at 4°C, resuspended in 50 µl of ice-cold PBS, and centrifuged once more at 550 × g for 5 minutes at 4 °C. The final cell pellets were resuspended in 50 µl of ice-cold lysis buffer (10 mM Tris-HCL, pH 7.4, 10 mM NaCl, 3 mM MgCl_2_, 0.1% IGEPAL CA-630) and immediately centrifuged at 550 × g for 10 minutes at 4 °C. The supernatant was discarded, cells were placed on wet ice, and the TDE1 Tagment DNA Enzyme and Buffer Kit (Illumina, #20034197) was utilized according to the manufacturer’s instructions. Briefly, chondrocytes were incubated in the TDE1 reaction buffer for 45 minutes at 37°C, followed by addition of 10 µl of 3 M sodium acetate to stop the reaction. Nucleic acid purification was performed using the Qiagen MinElute Kit (#28204) following the manufacturer’s instructions. Purified DNA was stored at −20°C until shipped to the CHOP Center for Spatial and Functional Genomics. Libraries were generated and sequenced in the same manner as indicated in the “hFOB, pediatric osteoblast, and osteoclast ATAC-seq library generation” methods section.

### ATAC-seq alignment and peak calling

ATAC-seq peaks were called for hFOBs, pediatric osteoblasts, osteoclasts, and chondrocytes using the ENCODE ATAC-seq pipeline^106^ and default settings. Briefly, this pipeline input pair-end reads from the biological replicates for each cell type and aligned them to GRCh38 using bowtie2^107^, removing any duplicate reads from the alignment. The pipeline then called narrow peaks independently for each replicate using macs2^108^ and removed peaks in ENCODE blacklist regions (ENCFF001TDO). Quality-control metrics were checked against the ENCODE recommended standards, and any data sets found not to meet standards, namely the 0, 8, and 12-day differentiated osteoclasts, were discarded from further analysis. For the S-LDSC analysis, we used the irreproducible discovery rate (IDR)^109^ optimal peak sets. For target selection using the hFOB and hMSC-osteoblast ATAC-seq results, we re-ran the ENCODE pipeline, aligning this time to hg19, and used the less stringent pooled peak sets.

The 4-day differentiated osteoclast peaks were also checked for overlap with the promoters of seven osteoclast marker genes: *CALCR*, *CA2*, *CTSK*, *MMP9*, *SPP1*, *ACP5*, *EDNRB*^101^. The promoters were defined as ±1kb from the GRCh38 transcription start sites as obtained from GeneCards^110^. All promoters except those for *CALCR* and *EDNRB* were overlapped by one or more peaks.

### GWAS summary statistics

Summary statistics for BMD (estimated by heel quantitative ultrasound) and bone fracture were obtained from the largest GWAS of each to date^8^. Each of these GWAS was executed on a population of white British individuals from the UK Biobank^111^ determined based on genetic similarity to the 1000 Genomes GBR subpopulation^102^. We also obtained GWAS summary statistics for 36 traits analyzed by the Pan UKBB project in an overlapping population of individuals based on genetic similarity to the 1000 Genomes EUR superpopulation^51,102^ (**Supplementary Table 18**). These metabolic and anthropometric phenotypes were manually selected to represent diverse areas of biology. All summary statistics were downloaded in hg19.

### Linkage disequilibrium score regression

We calculated genetic correlations and heritability enrichment in cell-type specific ATAC-seq and histone ChIP-seq peaks via cross-trait^112^ and stratified^36^ linkage disequilibrium (LD)-score regression^113^ respectively (v1.0.1; https://github.com/bulik/ldsc). We used the 1000 Genomes Project Phase III GRCh38 LD reference^114^ to calculate LD scores for the regressions and only retained SNPs included in the HapMap Project Phase 3 call set^115^. All GWAS summary statistics were lifted over to GRCh38 for the analyses using the UCSC LiftOver tool^116^ (https://genome.ucsc.edu/cgi-bin/hgLiftOver).

Stratified linkage disequilibrium score-regression (S-LDSC) heritability enrichments were calculated on top of the baseline LDSC model using 210 binary genomic annotations across 98 primary cell types and cell models. 198 of the annotations for 86 unique cell types were consolidated ChIP-seq narrow peak calls for four types of activating histone marks: H3K4me1, H3K4me3, H3K9ac, and H3K27ac. These peaks were obtained directly from the Roadmap Epigenomics Project^117,118^ and were supported by a minimum of 30M reads (**Supplementary Table 1**). These annotations were downloaded in hg19 and lifted over to GRCh38. We also included in the analysis the newly generated ATAC-seq peaks described above for pediatric hMSCs and hMSC-osteoblasts, chondrocytes, and RANKL-differentiated osteoclast models as well as reprocessed datasets obtained from the public domain. For the public domain datasets, we obtained raw FASTQ files for a H3K27ac ChIP-seq experiment in hFOBs^119^, a paired ATAC-seq / H3K27ac ChIP-seq experiment in RANKL-differentiated osteoclasts^120^, and two ATAC-seq studies previously published by our team for hMSC-osteoblasts^31^ and tonsillar-organoid sorted monocytes^43^. These FASTQs were reprocessed using standard ENCODE pipelines^106^ and aligned to GRCh38. The overlapping optimal peak sets were used for the reprocessed ChIP-seq datasets, and optimal IDR peaks were used for the reprocessed ATAC-seq experiments. The osteoclast peaks were also checked for overlap with the marker gene promoters. The ATAC-seq peaks overlapped all promoters except for *EDNRB*, and the H3K27ac ChIP-seq peaks overlapped 4 of 7 promoters (*CA2*, *CTSK*, *MMP9*, and *ACP5*).

Bonferroni-adjustment^121^ was used to correct for multiple testing across annotations in the S-LDSC analysis, and all annotations with an adjusted *P* < 0.05 were deemed to have significant heritability enrichment. Genetic correlations were adjusted for multiple testing via the Benjamini-Hochberg procedure^122^ as part of two separate analyses. In the first, genetic correlations were calculated for all traits with BMD (**Supplementary Table 19**), and in the second analysis, genetic correlations were calculated between all unique pairs of the weight and impedance related traits (**Supplementary Table 20**). Adjusted *P* < 0.05 were again considered significant for these analyses. Additionally, in the first analysis, two traits (hypoglycemia and type 2 diabetes) were found to have genetic correlations with BMD that could not be estimated by LDSC. These values were ignored when correcting for multiple testing and reporting results.

### hFOB promoter-focused Capture-C library preparation and sequencing

We followed the procedure previously published by our team for the generation and sequencing of the hFOB promoter-focused Capture-C libraries^123^. For this protocol, 10^7^ fixed hFOB cells were resuspended in dH2O supplemented with protease inhibitor cocktail and incubated on ice for 10 minutes twice. After setting aside 50 μl of cell suspension for pre-digestion QC, the remaining sample was divided into 6 tubes. All incubation reactions were carried out in a Thermomixer (BenchMark) shaking at 1,000 rpm. Samples were pre-digested for 1 hour at 37°C after adding 0.3% SDS, 1x NEB DpnII restriction enzyme buffer and dH2O. We then added a 1.7% solution of Triton X-100 to each tube and continued the incubation an additional hour. In the sample tubes only, we added 10 µL of DpnII (NEB, 50 U/μl) and continued the incubation until the end of the day when another 10 μl DpnII was added to each sample to digest overnight. The next morning, we added another 10 μl DpnII and incubated for a final 2−3 hours. We removed 100 µL of each digestion reaction, pooled them into two 1.5 ml tube, and set them aside for digestion efficiency QC. We heat inactivated the remaining samples at 65°C for 20 minutes, before cooling on ice for 20 minutes.

We ligated digested samples overnight at 16°C with T4 DNA ligase (HC ThermoFisher, 30 U/μl) and 1X ligase buffer. The next day, we spiked an additional T4 DNA ligase into each sample and incubated another few hours. We then de-crosslinked the samples overnight at 65°C with Proteinase K (20 mg/ml, Denville Scientific) along with pre-digestion and digestion controls. The following morning, we incubated the controls and ligated samples for 30 minutes. at 37°C with RNase A (Millipore) prior to phenol/chloroform extraction, ethanol precipitation at −20°C, and centrifugation at 4°C and 3000 rpm for 45 minutes to pellet the samples, while controls were pelleted at 14,000 rpm. Pellets were washed in 70% ethanol and centrifuged again as described above. We resuspended the 3C library and control pellets in dH2O and stored both at −20°C. We measured sample concentrations by Qubit and assessed digestion and ligation efficiencies by gel electrophoresis on a 0.9% agarose gel and by quantitative PCR (SYBR green, Thermo Fisher).

DNA from each 3C library (10 ug) was sheared to an average fragment size of 350bp using a QSonica Q800R (60% amplitude, 30 seconds on / 30 seconds off, 2-minute intervals for 5 total intervals) at 4°C. After shearing, DNA was purified using AMPureXP beads (Agencourt), DNA size was confirmed on a Bioanalyzer 2100 (Agilent) and DNA concentration measured via Qubit. Libraries were prepared for selection using the SureSelect XT Library Kit (Agilent) following the manufacturer protocol and once again bead purified, then checked for size and concentration as described above. One microgram of adaptor-ligated library was hybridized using the SureSelect XT capture kit (Agilent) and our custom-designed 41K promoter Capture-C probe set^31^. After amplification and purification, we assessed the quantity and quality of the captured libraries one final time. We paired-end sequenced all promoter-focused capture-C libraries on the Illumina NovaSeq 6000 platform with 51bp read length.

### Analysis of hFOB Capture-C data

We pre-processed paired-end reads from the three hFOB replicates using the HICUP pipeline^124^ (v0.5.9) aligning reads to the hg19 reference genome with bowtie2^107^. We called significant promoter interactions with genes from the GENCODE Release 19 gene set (GRCh37.p13)^125^ at 1-DpnII fragment resolution using CHiCAGO^126^ (v1.1.8) with default parameters except for binsize set to 2500. We also called significant interactions at 4-DpnII fragment resolution by artificially grouping four consecutive DpnII fragments and inputting them into CHiCAGO using default parameters except for “removeAdjacent” which was set to False. We considered interactions with a CHiCAGO score > 5 at either 1-fragment or 4-fragment resolution to be significant interactions and converted significant interactions to ibed format for use in variant to gene mapping.

### hFOB RNA-seq

Total RNA was isolated from hFOB cells using TRIzol reagent (Invitrogen) following manufacturer instructions, then purified using the Direct-zol RNA Plus Miniprep Kit (Zymol). After measuring concentration (Nanodrop, Invitrogen) and RNA integrity (RIN > 7, Bioanalyzer 2100, Agilent), RNA was depleted of rRNA using the QIAseq FastSelect RNA Removal Kit (Qiagen). RNA-seq libraries were prepared using the NEBNext Ultra II Directional RNA Library Prep Kit for Illumina (NEB) and NEBNext Multiplex Oligos for Illumina (Dual Index Primers, NEB) following standard protocols. Libraries were sequenced on an Illumina NovaSeq 6000, generating ~100 million paired-end 50 bp reads per sample. RNA-seq data were aligned to the hg19 genome with STAR v. 2.6.0c and gene counts were obtained using HTseq, with flags -f bam -r name -s reverse -t exon -m intersection-strict, to count genes from GENCODE Release 19 (GRCh37.p13) annotation plus annotation for lincRNAs and sno/miRNAs from the UCSC Table Browser (downloaded 7/7/2016). Normalized counts for the uniquely mapped read pairs were generated through the transcript per million read method with effective gene length and the resulting values were used in the computation of gene expression percentiles.

### CRISPRi Target Selection

We selected targets for the screen beginning with the list of 1,103 independent BMD signals reported by Morris *et al*.^8^ We identified LD-proxies for the signals at an r^2^ > 0.8 using SNiPA v3.4^127^ (https://snipa.org/snipa3/) and the 1000 Genomes Phase III hg19 LD reference^114^. 36 signals could not be mapped through SNiPA and were retained with only themselves as proxies. We mapped proxy variants to candidate effector genes in differentiated hFOBs and hMSC-osteoblasts by identifying promoter-interacting fragments that contained a proxy and overlapped an ATAC-seq peak in the same cell type. We discarded gene nominations for genes with expression of less than 1 transcript per million in the same matched cell types and any in which the implicating variant overlapped the promoter of an expressed gene. Having retained 88 signals with one or more linked gene, we found two pairs of these signals located within 1kb, the distance we assumed as the effective repressive range of CRISPRi^128^, and we combined them together. Additionally, we added three more targets we previously identified at genomic loci associated with pediatric bone density accrual^37^ bringing the total to 89 targets.

### Custom sgRNA pool target design

We designed synthetic guide RNAs (sgRNAs) for each target site using CRISPick^129,130^. We then input the CRISPick recommended sequences into FlashFry^131^ and discarded any candidate guides FlashFry flagged to have high GC content, polyT sequences, or multiple genomic targets. We iterated over the ranked list of remaining guides for each target and selected the top three whose binding sites did not overlap any of the previously selected guides. Positive and negative-control (scrambled) sgRNAs were designed and previously validated by Sigma. We selected the positive-control sgRNAs from an available list based on their expression in both hFOBs and HMC3 cells as measured by bulk RNA-seq. We considered joint expression as we were developing screens in both cell types that would be analyzed with a common analysis pipeline. The results of the HMC3 screen are not presented here, given that was for a separate unrelated trait.

### Generation of helper hFOBs and sgRNA configuration optimization

Helper hFOBs expressing the dCas9-CRISPRi-KRAB lentiviral construct under Blasticidin selection (10ug/ml,11 days) were first generated using the Sigma 10X CRISPRi Feature Barcode Optimization Kit (CRISPRI10X). Next, to test for the optimal sgRNA configuration, stocks of the helper hFOB-dCAS9-CRISPRi-KRAB cells were transduced with lentiviral pools containing one of four sgRNA capture sequence configurations and selected with Puromycin (1 ug/ml, 11 days): Capture Sequence One Stem (CS1-STEM), Capture Sequence One Three Prime (CS1-3’), Capture Sequence Two Stem (CS2-STEM), and Capture Sequence Two Three Prime (CS2-3’) (**Supplementary Fig. 24**). Each lentivirus pool contained sgRNA targeted to the *RAB1A* TSS and a Negative Control. Cells from all four configurations were subjected to 10X Genomics single cell analysis for both scRNA-seq (GEX library) and Feature Barcoded CRIPSR Capture (CRISPR Capture library). The optimal configuration was determined using both the best fraction of usable guide reads and the best −log_2_ fold change in *RAB1A* mRNA expression. The CS1-STEM configuration was determined to be optimal for hFOBs (**Supplementary Fig. 25**).

### Generation of CRISPRi sgRNA pool targeted hFOBs

To generate the hFOBs containing our custom sgRNAs, the same helper hFOB-dCas9-CRISPRi-KRAB cells were used for transduction. Cells were transduced at low MOI (0.2) plus polybrene (8 ug/ml) with a Sigma-Aldrich custom sgRNA lentiviral pool (titer = 5.3 × 10^8^ TU/ml). We selected an MOI of 0.2 to ensure that most viable cells would contain only one sgRNA and determined the optimal titer via the recommended procedure^132^. Under a Poisson model and perfect selection for transfected cells, ~90% of viable cells were expected to have one sgRNA. The lentiviral vectors followed Sigma-Aldrich’s standard CRISPRi-screen construct design (**Supplementary Fig. 26**) and were sequenced by the manufacturer to ensure quality prior to shipping (**Supplementary Table 2**). On day 2 post-transduction, cells were selected with Puromycin (1 ug/ml). Transduction was confirmed at day 8 by blue fluorescent protein (BFP) and frozen for stocks on day 11. Stock hFOB-CRISPRi-KRAB-Pooled-sgRNA cells were grown in 100mm plates under Blasticidin/Puromycin selection for 2 days at 33.5°C, then differentiated for 5 days at 39.5°C. Cells were removed from plates with TrypLE, counted, and diluted to 1000 cells/ul in DPBS+1% FBS. Viability was determined to be around 90% before 160,000 cells (8 lanes of 20K cells each) were processed for both 10X Genomics scRNA-seq (GEX libraries) and Feature Barcoded CRIPSR Capture (CRISPR Capture libraries) at the CHOP Center for Applied Genomics (CAG). Both sets of libraries were sequenced as eight pools on the Illumina Novaseq 6000 system using an S2-100 flow cell.

### Single-cell processing

Single-cell FASTQs were initially processed using the CellRanger pipeline (10X Genomics Cell Ranger 3.0.0)^133^ with default settings. We then used CellBender^134^ to denoise and filter the raw CellRanger outputs separately for each of the eight pools. The number of droplets and expected number of cells were visually estimated from each pool’s unique molecular identifier (UMI) curve, and we ran Cellbender with a learning rate of 0.00005, 150 training epochs, and a false-positive rate of 1%. After CellBender we used UMAP projections^54^ and violin plots implemented in Scanpy^135^ to visualize the remaining droplets for each pool and quickly recognized that all pools but the second contained two clusters of droplets, one with a high number of UMIs and genes per droplet and the other with a low level of cellular complexity. The second pool had an overall low level of complexity indicative of mostly empty droplets and was discarded. For the remaining pools we used the Leiden algorithm^136^ to cluster the cells and retained only the high complexity cluster. The remaining droplets were then combined across pools and visualized using UMAP and violin plots. We discarded droplets with >10% mitochondrial reads and >90,000 UMI to remove suspected dying cells and doublets respectively and retained 40,743 high-quality cells.

For comparisons of pseudo-bulk untargeted screen cells and bulk RNA-seq of osteoblast models, we extracted 2,340 cells that received only non-targeting guides and 7,703 that were either non-transfected cells or cells whose guides were not captured in the scRNA-seq. We calculated pseudo-bulk expression of each group in TPM and compared it to the bulk RNA-seq of hFOBs and hMSC-Osteoblasts used during target selection. We analyzed both principal components (PCs) and expression of osteoblast marker genes^46^.

In contrast for perturbation testing, we began with the 40,743 high-quality cells, and removed the 7,703 cells without detected sgRNAs. We then tested each non-targeting sgRNA for random assortment against each of the targeting guides using a Fisher Exact Test and a 2 × 2 contingency table. Results were corrected for multiple testing using the Benjamini-Hochberg procedure^122^ and a significance threshold of 0.05 was used. Finding 15 of 27 non-targeting guides preferentially assorted with one or more targeting guides, we discarded 5,655 remaining droplets containing more than one sgRNA, retaining 27,385 cells for the purposes of perturbation testing.

### Perturbation testing and visualizations

We next used the low-MOI version^50^ of SCEPTRE^49^ v0.3.0 (https://github.com/katsevich-lab/sceptre) to test all genes within 1Mb of each target site for perturbations. This method pools sgRNAs targeted to the same variants and tests them jointly using a permutation test comparing gene expression in cells receiving one of the sgRNAs targeting a particular site against cells receiving a non-targeting guide. Only cells with non-zero expression of the tested gene in each test are considered. We determined which genes to test for each target site by taking the full GENCODE Release 19 gene set (GRCh37.p13)^125^ and identifying all genes who overlapped or came within 1Mb of the binding sites for any of the sgRNAs for the target and were captured in the scRNA-seq. As covariate inputs into SCEPTRE, we included the number of UMIs per cell, the number of unique genes detected per cell, the pool in which each cell was sequenced, the mitochondrial read % per cell, and the top 15 gene expression PCs. We used Seurat v5.0.1^137^ to calculate the PCs via the recommended procedure on the 2,000 most variable genes determined via the vst method. We assumed targeted elements could be either enhancers or repressors and allowed for both possibilities by using a two-sided test. Statistical calibration was confirmed visually from the quantile-quantile plot. We corrected for multiple testing using the Benjamini-Hochberg procedure^122^. Adjusted *P* < 0.10 were considered significant.

We visualized genomic annotations at targets found to exhibit significant perturbations with pyGenomeTracks v3.8^138^ (https://pygenometracks.readthedocs.io/). Included among the tracks were the hg19-aligned hFOB ATAC-seq and Capture-C annotations described above as well as previously published ATAC-seq and Capture-C datasets for hMSC-osteoblasts^31^ and adipocytes differentiated from hMSCs (hMSC-adipocytes)^139^. For visualization purposes the basic GENCODE Release 19 gene set^125^ was used. Perturbed genes were plotted in red and all others in blue.

In a final use of SCEPTRE, we also tested osteoblast marker genes^46^ for trans-perturbations with the 20 siRNAs found to have one or more significant effects in the initial cis screen. We input into this test the same covariates except that we only needed the top 3 expression PCs to produce well calibrated results.

### Selection of gene targets for siRNA assays

Prior to beginning functional assays, we noted we had retained a few targets in our screen that, while not intersecting gene promoters, did overlap exons and were therefore not the focus of our work. We dropped the corresponding perturbed genes – *ADAT1*, *ADCY4*, and *FBXW4* – from consideration in our siRNA-based assays. For reference, the first two targets resided in occasionally-retained introns within the 5’-UTR of *ADAT1* and *ADCY4* respectively, and the last was a synonymous coding variant in *FBXW4*. We also dropped *RP11-242D8.1* from inclusion in the assays believing it was a false positive result since it was located at the same locus as the well-established BMD gene, *SOST*, and was the only gene to show increased expression in the screen, in contrast to the generally repressive nature of CRISPRi. However, after manually reviewing the sub-significant screen results for borderline genes with strong prior biological evidence of causality, we included *CALCRL* and *FAM118A*. *CALCRL* showed suggestive evidence of perturbation (*P* = 0.028) and is closely related to the calcitonin receptor that plays an essential role in bone biology^140,141^. *FAM118A* was borderline significant in the screen (*P* = 0.006) and supported by a chromatin loop observed in the hFOBs. This brought our total number of assayed genes to 21.

### hFOB siRNA treatments and alkaline phosphatase assay

Single cell suspensions of hFOBs were seeded into 24-well plates at 45-60K cells per well and allowed to adhere overnight. Transfections were carried out the next day using ON-TARGETplus SMARTpool siRNA purchased from Horizon Discovery (**Supplementary Table 25**) and Dharmafect-1 transfection reagent per the manufacturer’s protocol. Each SMARTpool consists of 4 siRNAs targeted to the same gene. The next day, growth media was replaced. The plate designated for differentiation into osteoblasts was placed at 39.5°C, while the permissive plate was kept at 33.5°C. Both plates were stained for ALP after 4 days using the Alkaline Phosphatase Staining Kit (Abcam, ab242286) following kit instructions. Plates were photographed and the images were split into 8-bit RGB images using Image J software. Images within the green channel were used to enumerate integrated density values within the cell culture area for each well as previously described^31^. Assays were repeated six times. Each replicate was conducted on a single plate. However, to align siRNAs between the hFOB and hMSC assays, for all plots, the hFOB results were split by siRNA into the two plate groups used to conduct the hMSC based assays. The control siRNA results were repeated under each plate to illustrate the tested comparisons between control and gene siRNAs.

### hMSC siRNA treatments and assays

Following our previously published protocol for conducting siRNA assays in hMSC models^31,37^, we obtained primary bone-marrow derived hMSCs isolated from healthy adult donors (**Supplementary Table 26**) and characterized them for cell surface expression (CD166 + CD90 + CD105+/CD36-CD34-CD10-CD11b-CD45-) and tri-lineage differentiation (osteoblastic, adipogenic, and chondrogenic) potential. We achieved experimental knockdown of candidate genes using siRNAs as in the hFOB cell models. For osteoblastic differentiation, we plated 15,000 cells/cm^2^ in alpha-MEM consisting of 16.5% FBS, 25 µg/ml Ascorbic acid-2-phosphate, 5 mM beta-glycerophosphate and 1% insulin-transferrin-selenous acid (osteogenic media) and stimulated them the next day with recombinant human BMP2 (300 ng/ml) (R&D Systems, MN) in serum-free osteogenic media. Cells were harvested at 72 hours following BMP2 treatment for alkaline phosphatase assessment and at 8–10 days for staining with Alizarin red S. ALP and ARS assay plates were scanned on a flatbed scanner and quantified by Image J after splitting the color images into 8-bit RGB images as described above.

For differentiation into hMSC-adipocytes, 30,000 cells were seeded on 24 well plates and transfected next day using Dharmafect-1. Cells were allowed to recover for 2 days and adipogenic differentiation was started using 10% FBS alpha-MEM supplemented with Indomethacin, IBMX, and Dexamethasone as described previously^31^. Media exchange was carried out every 3 days until staining with Oil Red O at 18-21 days. Lipid droplet accumulation was enumerated using Lionheart automated microscope in the Texas Red channel. 4X objective was used to take a montage of 25 different microscopic fields which were then stitched and quantified using the cell count feature. Representative images were taken with a 20x objective with DAPI nuclear staining for reference. Initially, each siRNA was tested in hMSCs from two biological donors. siRNAs exhibiting appreciable effects on Oil Red O staining and additionally showing significant effects on hMSC-osteoblast ALP and ARS were assayed in additional donors and tested for significance.

### qPCR validation and assaying of marker genes

Total RNA samples were prepared from differentiating cells using TRIzol^®^ reagent using standard procedure^142^. cDNA was synthesized using 600 ng of total RNA using High-Capacity cDNA Reverse Transcription Kit (Applied Biosystems) in a 20 μL reaction following our recently published procedure^142^. The resulting cDNA was diluted five times, and one microliter cDNA was amplified in a total PCR volume of 10 μL using Power SYBR Green PCR Master Mix (Applied Biosystems) and gene-specific primers in a QuantStudio Pro 6 (Applied Biosystems) following manufacturer’s recommendations. The sequences of primers except for those for *GAPDH*, *ID1*, *RUNX2*, *ALPL*, and *SP7* are provided in **Supplementary Table 27**. The primer sequences for those osteoblast marker genes are available in our previously published work^31^. Relative expression for each gene was normalized against *GAPDH* and expressed as fold change over control siRNA. Data from different donor lines were combined for reporting.

### DAPI staining procedure

ALP, ARS, and Oil Red O-stained assay plates were used to estimate relative cell numbers per well after primary data capture. Wells were washed three times with TBS-T (Tris buffered saline containing 0.1% Tween 20) for 20 minutes each and incubated with 200ng/ml DAPI in TBS-T at 4°C overnight in a shaking platform. Next day, the plates allowed to equilibrate to room temperature and further washed three times with TBS-T for 20 minutes each. Montage images were captured to cover ~70% of the well area using Lionheart FX automated microscope in the DAPI channel and stitched. Inbuilt cell count feature of Gene5 software was used to enumerate the number of cells in the stitched images. Relative cell percentage was calculated by normalizing cell numbers against undifferentiated cells from control siRNA transfected wells. Per-cell staining intensities were calculated at the well-level by dividing raw intensities by the DAPI cell counts. Additionally, the BMP2 differentiation is associated with significant cell death at the point at which the ARS assays were conducted (8-10 days post-differentiation) resulting in DAPI cell counts that do not tightly correlate with the cell number just before mineral deposition. Therefore, we used the undifferentiated wells in the ARS stain to estimate cell counts for both the control and BMP2-stimulated cells.

### Statistical testing of siRNA assays

The effects of siRNA knockdown on functional measures and cell counts were assessed by comparing wells with differentiated cells transduced with gene-targeting siRNAs to those targeted with scrambled, control siRNAs. Assays were designed to be at or near saturation in the unperturbed state, so one-sided paired Student’s t-tests were used to assess the loss of staining upon siRNA knockdown. For hMSC-based assays, where available, technical replicates reflecting multiple passages of donor lines were averaged together prior to using each donor measurement as an instance for statistical testing. ALP and ARS assays in hMSC-osteoblasts were both conducted in five donors, and adipogenesis assays were conducted in a minimum of 2 donors. If after 2 donors were tested for the adipogenesis assays, an siRNA was found to yield promising initial reductions in the number of intracellular lipid droplets and we detected significant responses for the siRNA in the osteoblast assays, we added more replicates and conducted significance testing. The decision to prioritize in this way was made due to the availability of matched stocks and the long differentiation time of the adipogenesis assays. We attempted to match the donors to the assays for the qPCR experiments, but some primers were found not to amplify in certain donors under control conditions and the results for those combinations of donors and genes were therefore dropped. Additionally, assay results for uncorrected staining intensities, cell counts, and per-cell staining intensities were also recalculated by normalizing all well values relative to the differentiated control-siRNA well on the corresponding plates. These plate-normalized results were reported in the supplement (**Supplementary Fig. 14**, **Supplementary Tables 12-14 and 16**).

### CAFEH multi-trait fine-mapping

We used the multi-trait fine-mapping algorithm, CAFEH (https://github.com/karltayeb/cafeh), to fine-map and colocalize shared causal BMD signals across the 38 GWAS described above. We first defined 501 BMD-relevant loci by adapting a previously reported approach^143^. This method involves tiling the genome into 250kb tiles and merging all adjacent tiles with one or more significant variants at a *P* < 10^−6^ threshold. Adjacent tiles were padded by 250kb on both sides to form loci and any overlapping loci were merged. Loci with one or more genome-wide significant variants (*P* < 5 × 10^−8^) were retained for analysis and the rest discarded. We then identified which non-BMD traits to fine-map at each locus by identifying any that had one or more genome-wide significant variants (*P* < 5 × 10^−8^) within the bounds of the BMD-defined locus. For each locus, we input the variants tested in all the mapped GWAS studies into CAFEH.

To execute signal fine-mapping, we downloaded published LD reference matrices calculated by the Pan-UKBB team in a subset of 421K participants from the UK Biobank who are genetically similar to individuals in the 1000 Genomes EUR superpopulation^51,102^. The UK Biobank-based populations used to create the LD matrices and conduct the GWAS heavily overlap and range in size from 347K to 427K individuals (**Supplementary Table 18**). We limited ourselves to these datasets, though larger GWAS are available for some of the traits, precisely so that we could minimize mismatch in the LD patterns within the GWAS study cohorts and between the GWAS cohorts and the individuals used to generate the LD matrices. Both types of mismatch are known to produce spurious results and inflated error rates^144–146^ that can be eliminated by conducting all the GWAS and generating the LD matrices in the exact same sample of individuals, a process known as “in-sample” fine-mapping. Our approach, limited by the summary-level data available in the public domain can be thought of as an approximation that leverages a strongly overlapping set of cohorts and reduces LD mismatch to the greatest extent possible.

We selected CAFEH for fine-mapping because among a limited number of multi-trait fine-mapping tools^53,147–149^, CAFEH appeared best suited to the specific task of efficiently mapping a variable selection of traits across a large number of BMD loci. We began applying CAFEH to each locus with the maximum number of signals set to the default, 10. If after initial mapping, at least one signal was detected with a purity value (the absolute correlation between the variants in the signal’s credible set) below 1%, we stopped mapping the locus. If, however, 10 signals were detected with purity values > 1%, we iteratively increased the number of signals by 1 and re-mapped the locus until at least one low-purity signal (< 1%) was detected or a maximum of 30 signals was reached.

Seeking to minimize suspicious results and prioritizing precision of reported signals over recall, we post-hoc filtered the CAFEH results by several metrics and reported only signals linked to BMD. First, we reported signals with purity > 50% as signals below this threshold represent instances where the model failed to distinguish between variants in moderately low LD. Second, we linked signals to traits only when the signals’ credible sets had a CAFEH activity score > 0.95 and one or more variants with *P* < 5 × 10^−8^ in the trait GWAS. These filters respectively capture signal-trait linkages with strong posterior evidence of trait causality under CAFEH’s Bayesian model and robust frequentist evidence of trait association. Lastly, we reported only signal-trait linkages where a variant in the credible set captures the maximum residual association for the corresponding signal in the given trait.

The residual association of each signal represents the remaining GWAS association at each SNP position after removing the effects of the other mapped signals. Mathematically, the residual association is the significance of the residual effect, written as ***r_−tk_*** for trait, *t*, and signal, *k*. The residual effect can be calculated by subtracting trait *t*’s first moments of the CAFEH joint model for all signals except *k*, from the GWAS βs (see equation 63 in CAFEH supplemental methods^53^) and then dividing the difference by the GWAS standard error. The residual effect approximately follows the unit normal distribution. Under a well-fit CAFEH model, plots of residual association for a given signal should appear similar to LocusZoom GWAS plots^150^ which either show a null distribution for traits not relevant to the signal or a uni-modal distribution with the credible set at or near the peak for traits linked confidently to the signal. However, in reviewing plots of these residual associations, we observed that often when variants in moderate LD appeared visually in the raw GWAS associations to be causal for different traits, CAFEH would group the variants into a single signal with the credible set driven by the trait with the stronger GWAS association. While these trait linkages could reflect true biology, we doubted that the model was correctly specified for the corresponding signals, so we removed them. Some of these trait linkages were removed by the activity score filter, and we removed the rest by ensuring that the credible set of each signal we reported lay right at the peak of the residual association for all the traits to which it was linked. For reference, we have provided a table of the number of BMD-linked signals detected under different filtering criteria (**Supplementary Table 28**) and a list of the 1,349 BMD-linked signals identified under the least stringent filtering criteria we considered (only a BMD activity score > 0.5) complete with the information to refilter the signals to any degree of stringency (**Supplementary Table 29**). After signal filtering, we hierarchically clustered the retained signals by their binarized trait linkages using the complete-linkage method with Euclidean distances implemented in pheatmap^151^ v1.0.12 (https://CRAN.R-project.org/package=pheatmap). For each signal, we also discretized the CAFEH weight means to −1, 0, and 1. These means reflect the effect-direction relationships between the traits linked to each signal. We standardized the discretized means so that the BMD weight mean would always have a value of 1, and used them to again cluster the signals hierarchically. Additionally, we visualized the binarized trait linkages using the umap package v0.2.10.0 (https://cran.r-project.org/web/packages/umap) and manually annotated the observed clusters by the dominant traits mapped to the signals found in each.

## DATA AVAILABILITY

All newly generated data sets are available on the Gene Expression Omnibus (GEO) at accession number GSE261284. Previously generated raw ATAC-seq and ChIP-seq read files for monocytes ^43^, osteoclasts ^120^, and hFOBs ^119^ are available on GEO at accessions GSE174658, GSE203587, and GSE152942 respectively. Raw capture-C, ATAC-seq, and RNA-seq reads from hMSC-Osteoblasts ^45^ are available on ArrayExpress with the following accession numbers: E-MTAB-6862, E-MTAB-6834, and E-MTAB-6835. GWAS summary statistics obtained from the Genetic Factors for Osteoporosis Consortium (GEFOS) and the Pan-UKBB team are available at the links provided in **Supplementary Table 18**.

## CODE AVAILABILITY

Public software packages are available at the citations and URLs listed. Custom code for this analysis has been deposited on GitHub (https://github.com/mconery/Grant_hFOB_CRISPRi).

## ACKNOWLEDGMENTS

The authors would like to thank AlloSource and the University of Colorado Interdisciplinary Joint Biology Program Biorepository for providing healthy human articular cartilage tissue samples for the chondrocyte ATAC-seq experiments and Ms. Samantha Landgrave for isolating chondrocytes from this tissue to support sequencing. Additionally, the authors would also like to acknowledge the Center for Applied Genomics (CAG) at CHOP for their assistance with the 10X scRNA-seq library generation and Dr. Winter Bruner for her help preparing samples for shipment.

## FUNDING

MJZ is supported by the University of Colorado Gates Grubstake Award; EK by the NSF (DMS 2113072 and DMS 2310654); ADW by NIAID (R01AI154773); ADW and SFAG by NIDDK (R01DK122586); BSZ by NCATS (UL1 TR001878); SFAG, BSZ, and KDH by the NICHD (R01 HD100406); SFAG and KDH by the NIA (R01 AG072705); SFAG and BV by the NIDDK (UM1 DK126194); KDH by the Henry Ruppenthal Family Professorship for Bioengineering and Orthopaedic Surgery; and SFAG by the Daniel B. Burke Endowed Chair for Diabetes Research.

## AUTHOR CONTRIBUTIONS

<u>Conceptualization</u>: MC, JAP, KT, ADW, BFV, AC, SFAG; <u>Methodology</u>: MC, JAP, YW, KT, MCP, EK, BFV, AC, SFAG; <u>Investigation</u>: MC, JAP, YW, KT, DAV, LJF; <u>Visualization</u>: MC, JAP, YW; <u>Funding acquisition</u>: MJZ, EK, ADW, BSZ, KDH, SFAG; <u>Supervision</u>: BFV, KDH, AC, SFAG; <u>Writing – original draft</u>: MC, JAP, YW, DAV; <u>Writing – review & editing</u>: MC, JAP, YW, KT, MCP, DAV, CLA, MJZ, EK, ADW, BSZ, BFV, KDH, AC, SFAG

## COMPETING INTERESTS

Authors declare that they have no competing interests.

## Notes

### Competing Interest Statement

The authors have declared no competing interest.

### Summary of Updates

We have added some minor analyses including a test for trans-perturbations of osteoblast marker genes and a more comprehensive examination of osteoblast marker expression in the screen hFOBs. We have also expanded the limitations section of the discussion.

## REFERENCES

1. Caliri, A., Filippis, L., Bagnato, G. & Bagnato, G. Osteoporotic fractures: Mortality and quality of life. Panminerva medica 49, 21–7 (2007).

2. Rizkallah, M. et al. Comparison of morbidity and mortality of hip and vertebral fragility fractures: Which one has the highest burden? Osteoporosis and Sarcopenia 6, 146–150 (2020).

3. Sattui, S. E. & Saag, K. G. Fracture mortality: associations with epidemiology and osteoporosis treatment. Nature Reviews Endocrinology 10, 592–602 (2014).

4. Krall, E. A. & Dawson-Hughes, B. Heritable and life-style determinants of bone mineral density. Journal of Bone and Mineral Research 8, 1–9 (1993).

5. Richards, J. B., Zheng, H.-F. & Spector, T. D. Genetics of osteoporosis from genome-wide association studies: advances and challenges. Nature Reviews Genetics 13, 576–588 (2012).

6. Ng, M. Y. M., Sham, P. C., Paterson, A. D., Chan, V. & Kung, A. W. C. Effect of Environmental Factors and Gender on the Heritability of Bone Mineral Density and Bone Size. Annals of Human Genetics 70, 428–438 (2006).

7. Kim, S. K. Identification of 613 new loci associated with heel bone mineral density and a polygenic risk score for bone mineral density, osteoporosis and fracture. PLOS ONE 13, e0200785 (2018).

8. Morris, J. A. et al. An atlas of genetic influences on osteoporosis in humans and mice. Nature Genetics 51, 258–266 (2019).

9. Johnell, O. et al. Predictive Value of BMD for Hip and Other Fractures. Journal of Bone and Mineral Research 20, 1185–1194 (2005).

10. Tak, Y. G. & Farnham, P. J. Making sense of GWAS: using epigenomics and genome engineering to understand the functional relevance of SNPs in non-coding regions of the human genome. Epigenetics & Chromatin 8, 57 (2015).

11. Sun, Q. et al. From GWAS variant to function: A study of ~148,000 variants for blood cell traits. Human Genetics and Genomics Advances 3, 100063 (2022).

12. Hindorff, L. A. et al. Potential etiologic and functional implications of genome-wide association loci for human diseases and traits. Proceedings of the National Academy of Sciences 106, 9362–9367 (2009).

13. Zhang, F. & Lupski, J. R. Non-coding genetic variants in human disease. Human Molecular Genetics 24, R102– R110 (2015).

14. Trynka, G. et al. Chromatin marks identify critical cell types for fine mapping complex trait variants. Nature Genetics 45, 124–130 (2013).

15. Degner, J. F. et al. DNase I sensitivity QTLs are a major determinant of human expression variation. Nature 482, 390–394 (2012).

16. Xie, S., Duan, J., Li, B., Zhou, P. & Hon, G. C. Multiplexed Engineering and Analysis of Combinatorial Enhancer Activity in Single Cells. Molecular Cell 66, 285–299.e5 (2017).

17. Xie, S., Armendariz, D., Zhou, P., Duan, J. & Hon, G. C. Global Analysis of Enhancer Targets Reveals Convergent Enhancer-Driven Regulatory Modules. Cell Reports 29, 2570–2578.e5 (2019).

18. Gasperini, M. et al. A Genome-wide Framework for Mapping Gene Regulation via Cellular Genetic Screens. Cell 176, 377–390.e19 (2019).

19. Alda-Catalinas, C. et al. Mapping the functional impact of non-coding regulatory elements in primary T cells through single-cell CRISPR screens. Genome Biology 25, 42 (2024).

20. Fulco, C. P. et al. Activity-by-contact model of enhancer–promoter regulation from thousands of CRISPR perturbations. Nature Genetics 51, 1664–1669 (2019).

21. Morris, J. A. et al. Discovery of target genes and pathways at GWAS loci by pooled single-cell CRISPR screens. Science 380, eadh7699 (2023).

22. Papalexi, E. et al. Characterizing the molecular regulation of inhibitory immune checkpoints with multimodal single-cell screens. Nature Genetics 53, 322–331 (2021).

23. Schraivogel, D. et al. Targeted Perturb-seq enables genome-scale genetic screens in single cells. Nature Methods 17, 629–635 (2020).

24. Shukla, A. & Huangfu, D. Decoding the noncoding genome via large-scale CRISPR screens. Current Opinion in Genetics & Development 52, 70–76 (2018).

25. Cooper, Y. A., Guo, Q. & Geschwind, D. H. Multiplexed functional genomic assays to decipher the noncoding genome. Human Molecular Genetics 31, R84–R96 (2022).

26. Wünnemann, F. et al. Multimodal CRISPR perturbations of GWAS loci associated with coronary artery disease in vascular endothelial cells. PLOS Genetics 19, e1010680 (2023).

27. Yihan Wang et al. Enhancer regulatory networks globally connect non-coding breast cancer loci to cancer genes. bioRxiv 2023.11.20.567880 (2023) doi:10.1101/2023.11.20.567880.

28. Armendariz, D. A. et al. CHD-associated enhancers shape human cardiomyocyte lineage commitment. eLife 12, e86206 (2023).

29. Wang, Z. et al. Landscape of enhancer disruption and functional screen in melanoma cells. Genome Biology 24, 248 (2023).

30. Yang, X. et al. Functional characterization of Alzheimer’s disease genetic variants in microglia. Nature Genetics 55, 1735–1744 (2023).

31. Chesi, A. et al. Genome-scale Capture C promoter interactions implicate effector genes at GWAS loci for bone mineral density. Nature Communications 10, 1260 (2019).

32. Calabrese, G. M. et al. Integrating GWAS and Co-expression Network Data Identifies Bone Mineral Density Genes SPTBN1 and MARK3 and an Osteoblast Functional Module. cels 4, 46–59.e4 (2017).

33. Guo, Y. et al. Integrating Epigenomic Elements and GWASs Identifies BDNF Gene Affecting Bone Mineral Density and Osteoporotic Fracture Risk. Scientific Reports 6, 30558 (2016).

34. Pippin, J. A., et al. CRISPR-Cas9–Mediated Genome Editing Confirms EPDR1 as an Effector Gene at the BMD GWAS-Implicated ‘STARD3NL’ Locus. JBMR Plus 5, e10531 (2021).

35. Dillard, L. J. et al. Single-Cell Transcriptomics of Bone Marrow Stromal Cells in Diversity Outbred Mice: A Model for Population-Level scRNA-Seq Studies. Journal of Bone and Mineral Research 38, 1350–1363 (2023).

36. Finucane, H. K. et al. Partitioning heritability by functional annotation using genome-wide association summary statistics. Nature Genetics 47, 1228–1235 (2015).

37. Cousminer, D. L. et al. Genome-wide association study implicates novel loci and reveals candidate effector genes for longitudinal pediatric bone accrual. Genome Biology 22, 1 (2021).

38. Medina-Gomez, C. et al. Bone mineral density loci specific to the skull portray potential pleiotropic effects on craniosynostosis. Communications Biology 6, 691 (2023).

39. Chen, D. et al. Osteogenic Differentiation Potential of Mesenchymal Stem Cells Using Single Cell Multiomic Analysis. Genes 14, (2023).

40. Abood, A. et al. Identification of Known and Novel Long Noncoding RNAs Potentially Responsible for the Effects of Bone Mineral Density (BMD) Genomewide Association Study (GWAS) Loci. Journal of Bone and Mineral Research 37, 1500–1510 (2022).

41. He, P. et al. Why SNP rs3755955 is associated with human bone mineral density? A molecular and cellular study in bone cells. Molecular and Cellular Biochemistry 477, 455–468 (2022).

42. Xia, Q. et al. The type 2 diabetes presumed causal variant within TCF7L2 resides in an element that controls the expression of ACSL5. Diabetologia 59, 2360–2368 (2016).

43. Pahl, M. C. et al. Implicating effector genes at COVID-19 GWAS loci using promoter-focused Capture-C in disease-relevant immune cell types. Genome Biology 23, 125 (2022).

44. Palermo, J. et al. Variant-to-gene mapping followed by cross-species genetic screening identifies GPI-anchor biosynthesis as a regulator of sleep. Science Advances 9, eabq0844 (2023).

45. Su, C. et al. Mapping effector genes at lupus GWAS loci using promoter Capture-C in follicular helper T cells. Nature Communications 11, 3294 (2020).

46. Sojan, J. M. et al. Bacillus subtilis Modulated the Expression of Osteogenic Markers in a Human Osteoblast Cell Line. Cells 12, (2023).

47. Freiholtz, D. et al. SPP1/osteopontin: a driver of fibrosis and inflammation in degenerative ascending aortic aneurysm? Journal of Molecular Medicine 101, 1323–1333 (2023).

48. Lamort, A.-S., Giopanou, I., Psallidas, I. & Stathopoulos, G. T. Osteopontin as a Link between Inflammation and Cancer: The Thorax in the Spotlight. Cells 8, (2019).

49. Barry, T., Wang, X., Morris, J. A., Roeder, K. & Katsevich, E. SCEPTRE improves calibration and sensitivity in single-cell CRISPR screen analysis. Genome Biology 22, 344 (2021).

50. Timothy Barry, Kaishu Mason, Kathryn Roeder, & Eugene Katsevich. Robust differential expression testing for single-cell CRISPR screens at low multiplicity of infection. bioRxiv 2023.05.15.540875 (2023) doi:10.1101/2023.05.15.540875.

51. The Pan UKBB Team. Pan UKBB. https://pan.ukbb.broadinstitute.org.

52. Qu, Y. et al. Genetic Correlation, Shared Loci, and Causal Association Between Sex Hormone-Binding Globulin and Bone Mineral Density: Insights From a Large-Scale Genomewide Cross-Trait Analysis. Journal of Bone and Mineral Research 38, 1635–1644 (2023).

53. Arvanitis, M., Tayeb, K., Strober, B. J. & Battle, A. Redefining tissue specificity of genetic regulation of gene expression in the presence of allelic heterogeneity. The American Journal of Human Genetics 109, 223–239 (2022).

54. McInnes, L., Healy, J. & Melville, J. Umap: Uniform manifold approximation and projection for dimension reduction. *arXiv preprint arXiv:1802.03426* (2018).

55. Sheu, Y.-T. et al. Nuclear Receptor Coactivator-3 Alleles Are Associated with Serum Bioavailable Testosterone, Insulin-Like Growth Factor-1, and Vertebral Bone Mass in Men. The Journal of Clinical Endocrinology & Metabolism 91, 307–312 (2006).

56. Patel, M. S. et al. Alleles of the Estrogen Receptor α-Gene and an Estrogen Receptor Cotranscriptional Activator Gene, Amplified in Breast Cancer-1 (AIB1), Are Associated with Quantitative Calcaneal Ultrasound. Journal of Bone and Mineral Research 15, 2231–2239 (2000).

57. Salazar-Silva, R. et al. NCOA3 identified as a new candidate to explain autosomal dominant progressive hearing loss. Human Molecular Genetics 29, 3691–3705 (2020).

58. Schmahl, J., Raymond, C. S. & Soriano, P. PDGF signaling specificity is mediated through multiple immediate early genes. Nature genetics 39, 52–60 (2007).

59. Lahoud, M. H. et al. Gene targeting of Desrt, a novel ARID class DNA-binding protein, causes growth retardation and abnormal development of reproductive organs. Genome research 11, 1327–1334 (2001).

60. Whitson, R. H., Tsark, W., Huang, T. H. & Itakura, K. Neonatal mortality and leanness in mice lacking the ARID transcription factor Mrf-2. Biochemical and biophysical research communications 312, 997–1004 (2003).

61. Hata, K. et al. Arid5b facilitates chondrogenesis by recruiting the histone demethylase Phf2 to Sox9-regulated genes. Nature Communications 4, 2850 (2013).

62. Amano, K. et al. Arid5a cooperates with Sox9 to stimulate chondrocyte-specific transcription. MBoC 22, 1300–1311 (2011).

63. Shen, Y. et al. MicroRNA-877-5p promotes osteoblast differentiation by targeting EIF4G2 expression. Journal of Orthopaedic Surgery and Research 19, 134 (2024).

64. Wang, Z., Ding, X., Cao, F., Zhang, X. & Wu, J. Bone mesenchymal stem cells promote extracellular matrix remodeling of degenerated nucleus pulposus cells via the miR-101-3p/EIF4G2 axis. Frontiers in Bioengineering and Biotechnology 9, 642502 (2021).

65. Gao, S., et al. MicroRNA-197 regulates chondrocyte proliferation, migration, and inflammation in pathogenesis of osteoarthritis by targeting EIF4G2. Bioscience Reports 40, BSR20192095 (2020).

66. Xie, X. et al. MicroRNA-379 inhibits the proliferation, migration and invasion of human osteosarcoma cells by targetting EIF4G2. Bioscience reports 37, BSR20160542 (2017).

67. Kim, K. et al. Transcriptional Expression in Human Periodontal Ligament Cells Subjected to Orthodontic Force: An RNA-Sequencing Study. Journal of Clinical Medicine 9, (2020).

68. Deshar, R., Cho, E.-B., Yoon, S. K. & Yoon, J.-B. CC2D1A and CC2D1B regulate degradation and signaling of EGFR and TLR4. Biochemical and Biophysical Research Communications 480, 280–287 (2016).

69. Ventimiglia, L. N. et al. CC2D1B coordinates ESCRT-III activity during the mitotic reformation of the nuclear envelope. Developmental cell 47, 547–563 (2018).

70. Martinelli, N. et al. CC2D1A Is a Regulator of ESCRT-III CHMP4B. Journal of Molecular Biology 419, 75–88 (2012).

71. Li, C. Exploring the Cellular Roles of CC2D1A/B. PQDT - Global (The University of Manchester (United Kingdom), England, 2019).

72. Amariuta, T., Siewert-Rocks, K. & Price, A. L. Modeling tissue co-regulation estimates tissue-specific contributions to disease. Nature Genetics 55, 1503–1511 (2023).

73. Ongen, H. et al. Estimating the causal tissues for complex traits and diseases. Nature Genetics 49, 1676–1683 (2017).

74. Benjamin J. Strober, Martin Jinye Zhang, Tiffany Amariuta, Jordan Rossen, & Alkes L. Price. Fine-mapping causal tissues and genes at disease-associated loci. medRxiv 2023.11.01.23297909 (2023) doi:10.1101/2023.11.01.23297909.

75. Mostafavi, H., Spence, J. P., Naqvi, S. & Pritchard, J. K. Systematic differences in discovery of genetic effects on gene expression and complex traits. Nature Genetics 55, 1866–1875 (2023).

76. Grundberg, E. et al. Population genomics in a disease targeted primary cell model. Genome Res. 19, 1942– 1952 (2009).

77. Mullin, B. H. et al. Expression Quantitative Trait Locus Study of Bone Mineral Density GWAS Variants in Human Osteoclasts. Journal of Bone and Mineral Research 33, 1044–1051 (2018).

78. Magnusson, P., Degerblad, M., Sääf, M., Larsson, L. & Thorén, M. Different Responses of Bone Alkaline Phosphatase Isoforms During Recombinant Insulin-like Growth Factor-I (IGF-I) and During Growth Hormone Therapy in Adults with Growth Hormone Deficiency. Journal of Bone and Mineral Research 12, 210–220 (1997).

79. Bevier, W. C. et al. Relationship of body composition, muscle strength, and aerobic capacity to bone mineral density in older men and women. Journal of Bone and Mineral Research 4, 421–432 (1989).

80. Sutter, T. et al. Relationships between muscle mass, strength and regional bone mineral density in young men. PLOS ONE 14, e0213681 (2019).

81. Snow-Harter, C., Whalen, R., Myburgh, K., Arnaud, S. & Marcus, R. Bone mineral density, muscle strength, and recreational exercise in men. Journal of Bone and Mineral Research 7, 1291–1296 (1992).

82. Henderson, N. K., Price, R. I., Cole, J. H., Gutteridge, D. H. & Bhagat, C. I. Bone density in young women is associated with body weight and muscle strength but not dietary intakes. Journal of Bone and Mineral Research 10, 384–393 (1995).

83. Ho-Pham, L. T., Nguyen, U. D. T. & Nguyen, T. V. Association Between Lean Mass, Fat Mass, and Bone Mineral Density: A Meta-analysis. The Journal of Clinical Endocrinology & Metabolism 99, 30–38 (2014).

84. Katzmarzyk, P. T. et al. Relationship between abdominal fat and bone mineral density in white and African American adults. Bone 50, 576–579 (2012).

85. Kim, W. et al. The relationship between body fat and bone mineral density in Korean men and women. J Bone Miner Metab 32, 709–717 (2014).

86. Lee, S. J., Lee, J.-Y. & Sung, J. Obesity and Bone Health Revisited: A Mendelian Randomization Study for Koreans. Journal of Bone and Mineral Research 34, 1058–1067 (2019).

87. Song, J. et al. Causal associations of hand grip strength with bone mineral density and fracture risk: A mendelian randomization study. Frontiers in Endocrinology 13, (2022).

88. Liu, C. et al. Osteoporosis and sarcopenia-related traits: A bi-directional Mendelian randomization study. Frontiers in Endocrinology 13, (2022).

89. Felson, D. T., Zhang, Y., Hannan, M. T. & Anderson, J. J. Effects of weight and body mass index on bone mineral density in men and women: The framingham study. Journal of Bone and Mineral Research 8, 567– 573 (1993).

90. Ma, B. et al. Causal Associations of Anthropometric Measurements With Fracture Risk and Bone Mineral Density: A Mendelian Randomization Study. Journal of Bone and Mineral Research 36, 1281–1287 (2021).

91. Zhu, K. et al. Relationship between visceral adipose tissue and bone mineral density in Australian baby boomers. Osteoporos Int 31, 2439–2448 (2020).

92. Bland, V. L. et al. Metabolically favorable adiposity and bone mineral density: a Mendelian randomization analysis. Obesity 31, 267–278 (2023).

93. Hu, J. et al. Associations of visceral adipose tissue with bone mineral density and fracture: observational and Mendelian randomization studies. Nutrition & Metabolism 19, 45 (2022).

94. Liu, P.-Y., Ilich, J. Z., Brummel-Smith, K. & Ghosh, S. New Insight into Fat, Muscle and Bone Relationship in Women: Determining the Threshold at Which Body Fat Assumes Negative Relationship with Bone Mineral Density. Int J Prev Med 5, 1452–1463 (2014).

95. Kemp, J. P., Sayers, A., Smith, G. D., Tobias, J. H. & Evans, D. M. Using Mendelian randomization to investigate a possible causal relationship between adiposity and increased bone mineral density at different skeletal sites in children. International Journal of Epidemiology 45, 1560–1572 (2016).

96. Mullin, B. H. et al. Characterisation of genetic regulatory effects for osteoporosis risk variants in human osteoclasts. Genome Biology 21, 80 (2020).

97. He, D. et al. A longitudinal genome-wide association study of bone mineral density mean and variability in the UK Biobank. Osteoporosis International 34, 1907–1916 (2023).

98. Dong, H. et al. Comprehensive Analysis of the Genetic and Epigenetic Mechanisms of Osteoporosis and Bone Mineral Density. Frontiers in Cell and Developmental Biology 8, (2020).

99. Timshel, P. N., Thompson, J. J. & Pers, T. H. Genetic mapping of etiologic brain cell types for obesity. eLife 9, e55851 (2020).

100. Greenbaum, J. et al. A multiethnic whole genome sequencing study to identify novel loci for bone mineral density. Human Molecular Genetics 31, 1067–1081 (2022).

101. Takeshita, S., Kaji, K. & Kudo, A. Identification and Characterization of the New Osteoclast Progenitor with Macrophage Phenotypes Being Able to Differentiate into Mature Osteoclasts. Journal of Bone and Mineral Research 15, 1477–1488 (2000).

102. Fairley, S., Lowy-Gallego, E., Perry, E. & Flicek, P. The International Genome Sample Resource (IGSR) collection of open human genomic variation resources. Nucleic Acids Research 48, D941–D947 (2020).

103. Cody, J. J. et al. A simplified method for the generation of human osteoclasts in vitro. Int J Biochem Mol Biol 2, 183–189 (2011).

104. Susa, M., Luong-Nguyen, N.-H., Cappellen, D., Zamurovic, N. & Gamse, R. Human primary osteoclasts: in vitro generation and applications as pharmacological and clinical assay. J Transl Med 2, 6 (2004).

105. Shen, J. et al. DNA methyltransferase 3b regulates articular cartilage homeostasis by altering metabolism. JCI Insight 2, e93612.

106. Benjamin C. Hitz et al. The ENCODE Uniform Analysis Pipelines. bioRxiv 2023.04.04.535623 (2023) doi:10.1101/2023.04.04.535623.

107. Langmead, B. & Salzberg, S. L. Fast gapped-read alignment with Bowtie 2. Nature Methods 9, 357–359 (2012).

108. John M. Gaspar. Improved peak-calling with MACS2. bioRxiv 496521 (2018) doi:10.1101/496521.

109. Qunhua Li, James B. Brown, Haiyan Huang, & Peter J. Bickel. Measuring reproducibility of high-throughput experiments. The Annals of Applied Statistics 5, 1752–1779 (2011).

110. Stelzer, G. et al. The GeneCards Suite: From Gene Data Mining to Disease Genome Sequence Analyses. Current Protocols in Bioinformatics 54, 1.30.1–1.30.33 (2016).

111. Sudlow, C. et al. UK Biobank: An Open Access Resource for Identifying the Causes of a Wide Range of Complex Diseases of Middle and Old Age. PLOS Medicine 12, e1001779 (2015).

112. Bulik-Sullivan, B. et al. An atlas of genetic correlations across human diseases and traits. Nature Genetics 47, 1236–1241 (2015).

113. Bulik-Sullivan, B. K. et al. LD Score regression distinguishes confounding from polygenicity in genome-wide association studies. Nature Genetics 47, 291–295 (2015).

114. Auton, A. et al. A global reference for human genetic variation. Nature 526, 68–74 (2015).

115. Altshuler, D. M. et al. Integrating common and rare genetic variation in diverse human populations. Nature 467, 52–58 (2010).

116. Hinrichs, A. S. et al. The UCSC Genome Browser Database: update 2006. Nucleic Acids Research 34, D590– D598 (2006).

117. Bernstein, B. E. et al. The NIH Roadmap Epigenomics Mapping Consortium. Nature Biotechnology 28, 1045– 1048 (2010).

118. Kundaje, A. et al. Integrative analysis of 111 reference human epigenomes. Nature 518, 317–30 (2015).

119. Cottone, L. et al. Aberrant paracrine signalling for bone remodelling underlies the mutant histone-driven giant cell tumour of bone. Cell Death & Differentiation 29, 2459–2471 (2022).

120. Bae, S. et al. RANKL-responsive epigenetic mechanism reprograms macrophages into bone-resorbing osteoclasts. Cellular & Molecular Immunology 20, 94–109 (2023).

121. Bonferroni, C. Teoria statistica delle classi e calcolo delle probabilita. Pubblicazioni del R Istituto Superiore di Scienze Economiche e Commericiali di Firenze 8, 3–62 (1936).

122. Benjamini, Y. & Hochberg, Y. Controlling the False Discovery Rate: A Practical and Powerful Approach to Multiple Testing. Journal of the Royal Statistical Society: Series B (Methodological) 57, 289–300 (1995).

123. Su, C. et al. 3D promoter architecture re-organization during iPSC-derived neuronal cell differentiation implicates target genes for neurodevelopmental disorders. Progress in Neurobiology 201, 102000 (2021).

124. Wingett, S. W., et al. HiCUP: pipeline for mapping and processing Hi-C data. Preprint at 10.12688/f1000research.7334.1 (2015).

125. Frankish, A. et al. GENCODE 2021. Nucleic Acids Research 49, D916–D923 (2021).

126. Cairns, J., et al. CHiCAGO: robust detection of DNA looping interactions in Capture Hi-C data. Genome Biology 17, 127 (2016).

127. Arnold, M., Raffler, J., Pfeufer, A., Suhre, K. & Kastenmüller, G. SNiPA: an interactive, genetic variant-centered annotation browser. Bioinformatics 31, 1334–1336 (2015).

128. Thakore, P. I. et al. Highly specific epigenome editing by CRISPR-Cas9 repressors for silencing of distal regulatory elements. Nature Methods 12, 1143–1149 (2015).

129. Doench, J. G. et al. Optimized sgRNA design to maximize activity and minimize off-target effects of CRISPR-Cas9. Nature Biotechnology 34, 184–191 (2016).

130. Sanson, K. R. et al. Optimized libraries for CRISPR-Cas9 genetic screens with multiple modalities. Nature Communications 9, 5416 (2018).

131. McKenna, A. & Shendure, J. FlashFry: a fast and flexible tool for large-scale CRISPR target design. BMC Biology 16, 74 (2018).

132. Sigma Aldrich. CRISPRi Human Whole Genome and Long Non-Coding Library Screening User Manual. (2021).

133. Zheng, G. X. Y. et al. Massively parallel digital transcriptional profiling of single cells. Nature Communications 8, 14049 (2017).

134. Fleming, S. J. et al. Unsupervised removal of systematic background noise from droplet-based single-cell experiments using CellBender. Nature Methods 20, 1323–1335 (2023).

135. Wolf, F. A., Angerer, P. & Theis, F. J. SCANPY: large-scale single-cell gene expression data analysis. Genome Biology 19, 15 (2018).

136. Traag, V. A., Waltman, L. & van Eck, N. J. From Louvain to Leiden: guaranteeing well-connected communities. Scientific Reports 9, 5233 (2019).

137. Hao, Y. et al. Dictionary learning for integrative, multimodal and scalable single-cell analysis. Nature Biotechnology (2023) doi:10.1038/s41587-023-01767-y.

138. Lopez-Delisle, L. et al. pyGenomeTracks: reproducible plots for multivariate genomic datasets. Bioinformatics 37, 422–423 (2021).

139. Hammond, R. K. et al. Biological constraints on GWAS SNPs at suggestive significance thresholds reveal additional BMI loci. eLife 10, e62206 (2021).

140. Xie, J. et al. Calcitonin and Bone Physiology: In Vitro, In Vivo, and Clinical Investigations. International Journal of Endocrinology 2020, 3236828 (2020).

141. Naot, D., Musson, D. S. & Cornish, J. The Activity of Peptides of the Calcitonin Family in Bone. Physiological Reviews 99, 781–805 (2019).

142. Kaur, G., et al. Osteoporosis GWAS-implicated DNM3 locus contextually regulates osteoblastic and chondrogenic fate of mesenchymal stem/progenitor cells through oscillating miR-199a-5p levels. JBMR Plus 8, ziae051 (2024).

143. Anurag Verma et al. Diversity and Scale: Genetic Architecture of 2,068 Traits in the VA Million Veteran Program. medRxiv 2023.06.28.23291975 (2023) doi:10.1101/2023.06.28.23291975.

144. Chen, W. et al. Improved analyses of GWAS summary statistics by reducing data heterogeneity and errors. Nature Communications 12, 7117 (2021).

145. Kanai, M. et al. Meta-analysis fine-mapping is often miscalibrated at single-variant resolution. Cell genomics 2, (2022).

146. Yang, Z. et al. CARMA is a new Bayesian model for fine-mapping in genome-wide association meta-analyses. Nature Genetics 55, 1057–1065 (2023).

147. Wallace, C. A more accurate method for colocalisation analysis allowing for multiple causal variants. PLOS Genetics 17, e1009440 (2021).

148. Zhou, F. et al. Leveraging information between multiple population groups and traits improves fine-mapping resolution. Nature Communications 14, 7279 (2023).

149. Yuxin Zou, Peter Carbonetto, Dongyue Xie, Gao Wang, & Matthew Stephens. Fast and flexible joint fine-mapping of multiple traits via the Sum of Single Effects model. bioRxiv 2023.04.14.536893 (2024) doi:10.1101/2023.04.14.536893.

150. Pruim, R. J. et al. LocusZoom: regional visualization of genome-wide association scan results. Bioinformatics 26, 2336–2337 (2010).

151. Kolde, R. & Kolde, M. R. Package ‘pheatmap’. R package 1, 790 (2015).

