## Supplementary Figures for "GWAS-Informed data integration and non-coding CRISPRi screen illuminate genetic etiology of bone mineral density"

Mitchell Conery *et al.*

### Supplementary Figures

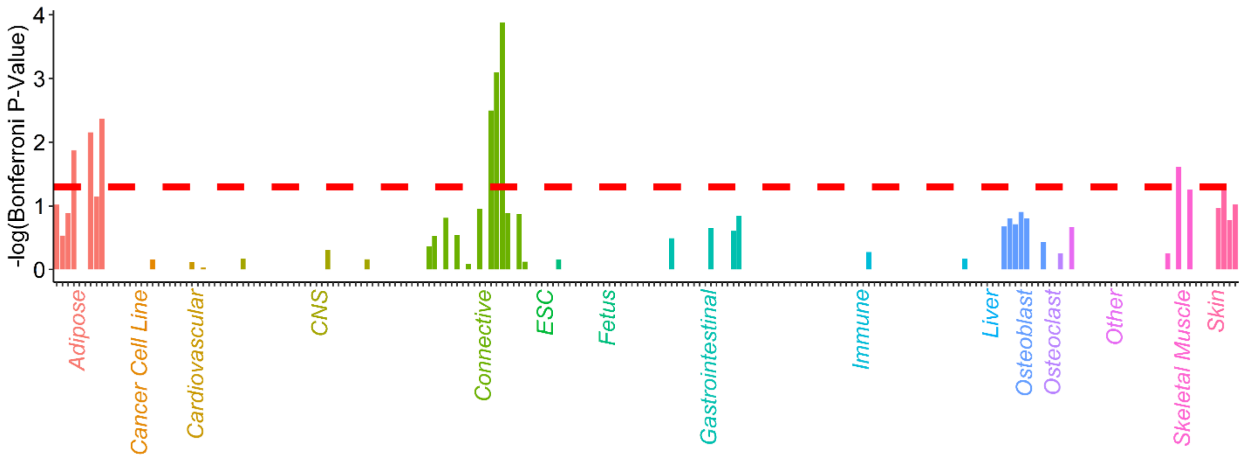


**Supplementary Fig. 1 - Bone Fracture Partitioned Heritability Enrichments across 98 Cell Types.**

Each bar in the figure represents a particular genomic annotation (H3K27ac, H3K9ac, H3K4me1, H3K4me3, or open-chromatin) measured in a specific primary cell type or cell model. Negative log10 Bonferroni-adjusted p-values are plotted along the y-axis. The dashed line reflects a Bonferroni-adjusted significance cutoff of 0.05. Coloring reflects manually curated tissue categories for each cell type.

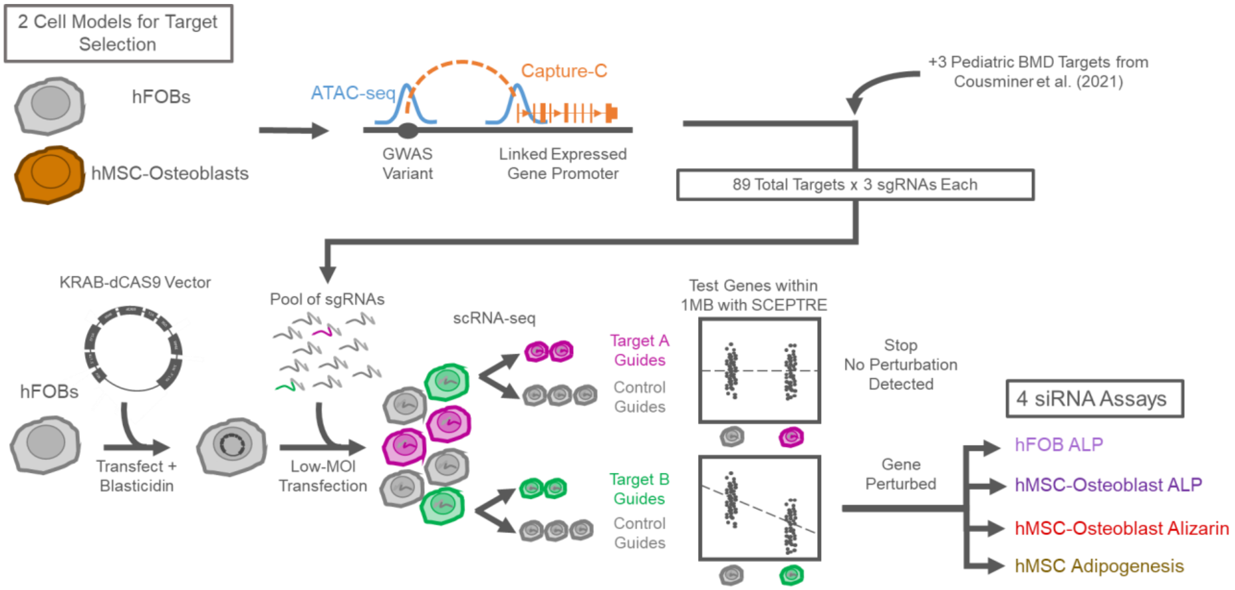


**Supplementary Fig. 2 - Design of Target-Selection, hFOB CRISPRi Screen, and siRNA Functional Assays.**

Schematic overview of main functional experiments.

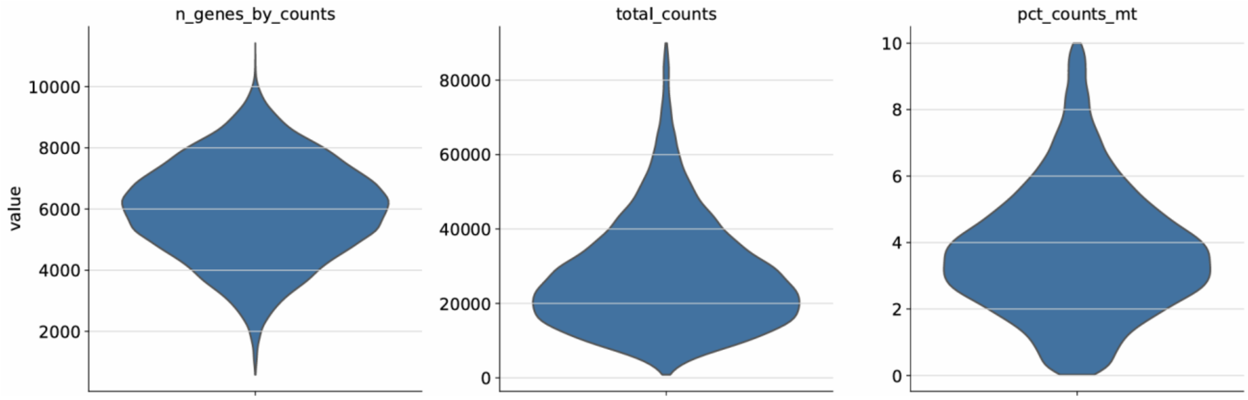


**Supplementary Fig. 3 - Distributions of Quality Control Metrics in 27,385 Cells Retained after QC Filters.**

Violin plots showing the number of unique genes per droplet, the number of UMIs per droplet, and the percent of reads that map to the mitochondrial genome in each droplet.

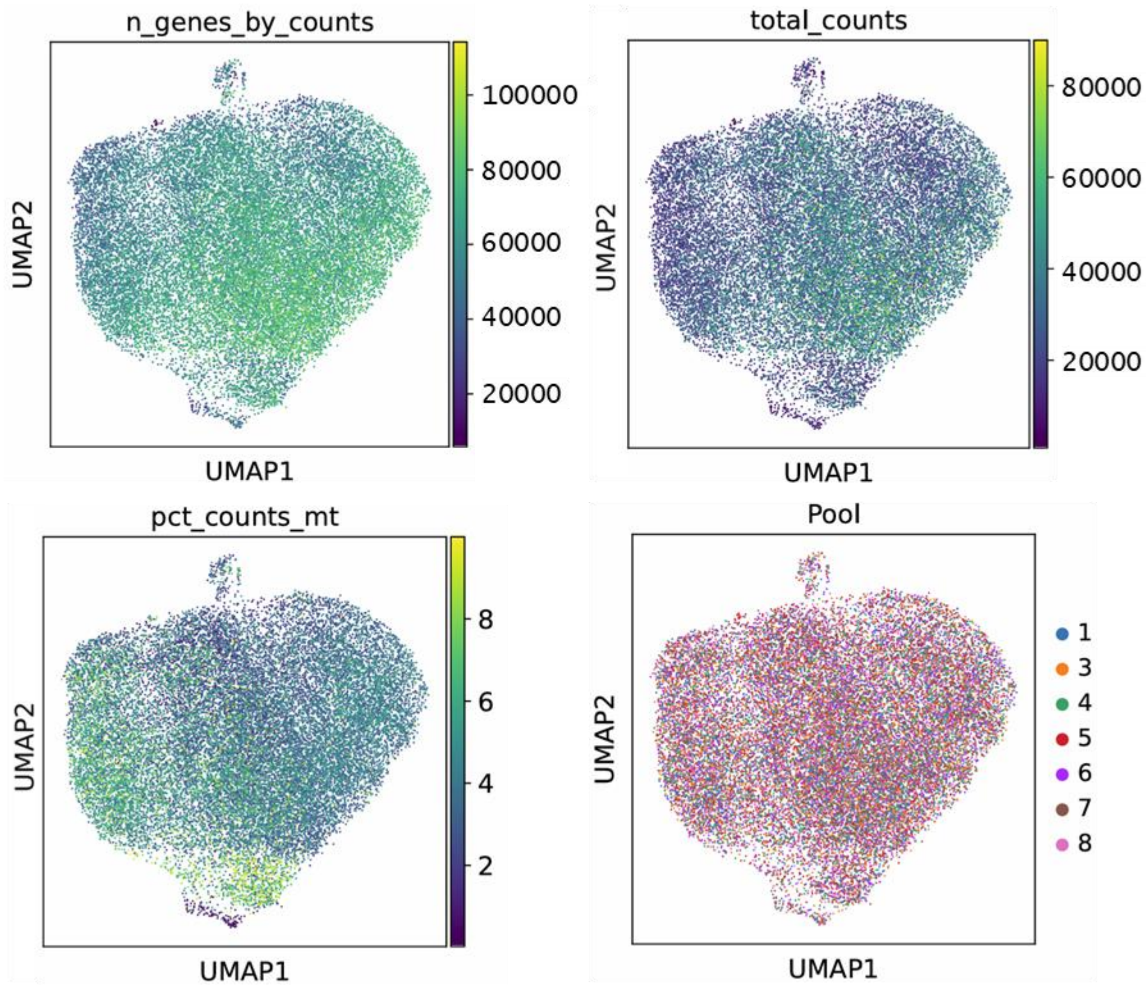


**Supplementary Fig. 4 - UMAP Embeddings of 27,385 Cells Passing Quality-Control Filters.**

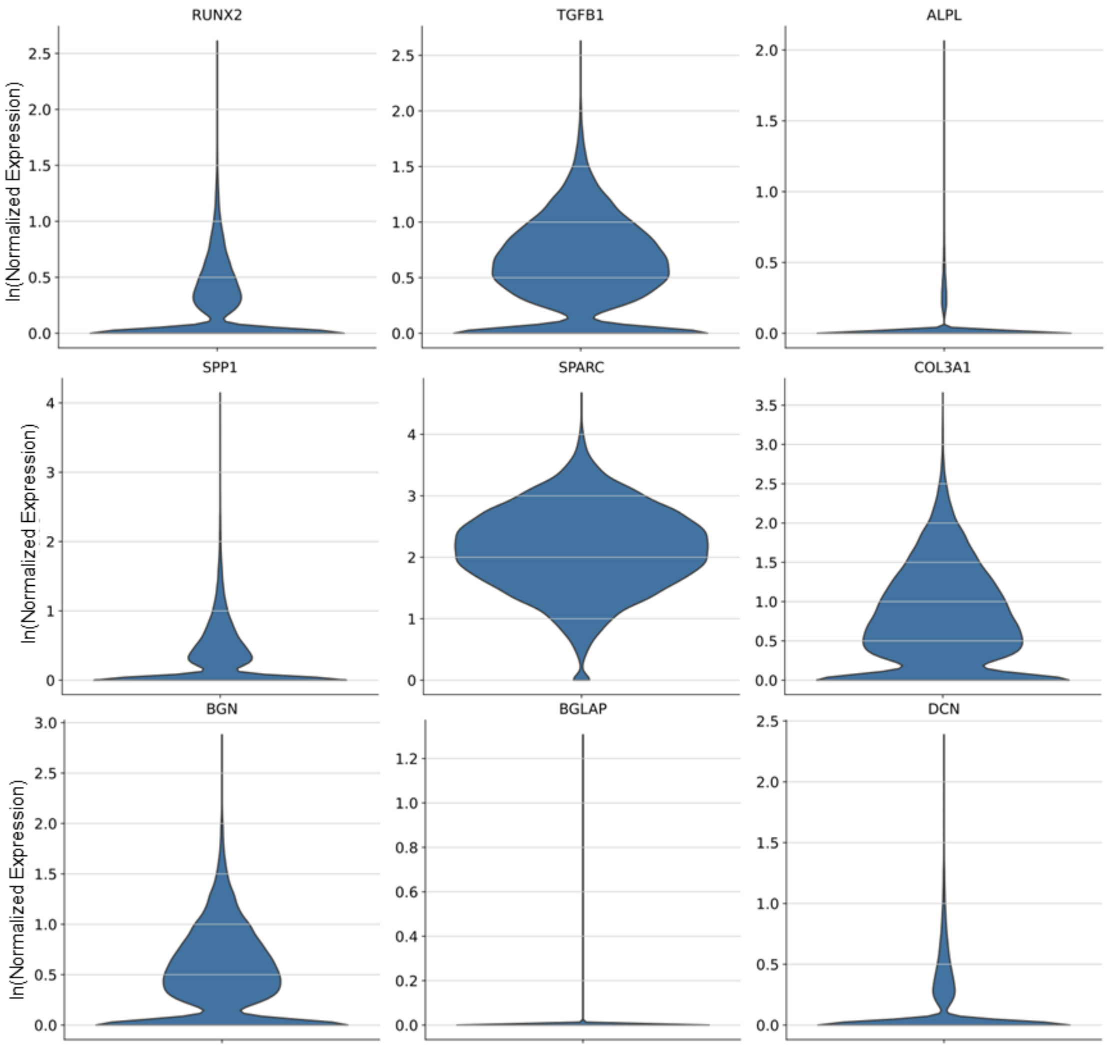


**Supplementary Fig. 5 - Osteoblast Marker Gene Expression in 27,385 Cells Retained after QC Filters**

Violin plots of osteoblast proliferation markers (top row), maturation markers (middle row), and mineralization markers (bottom row).

**A**

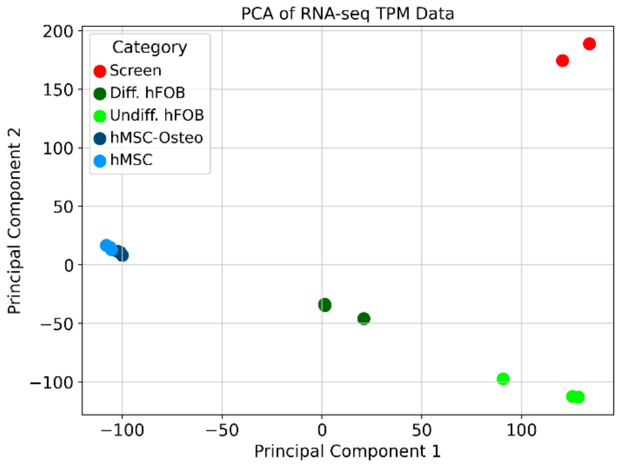


**B**

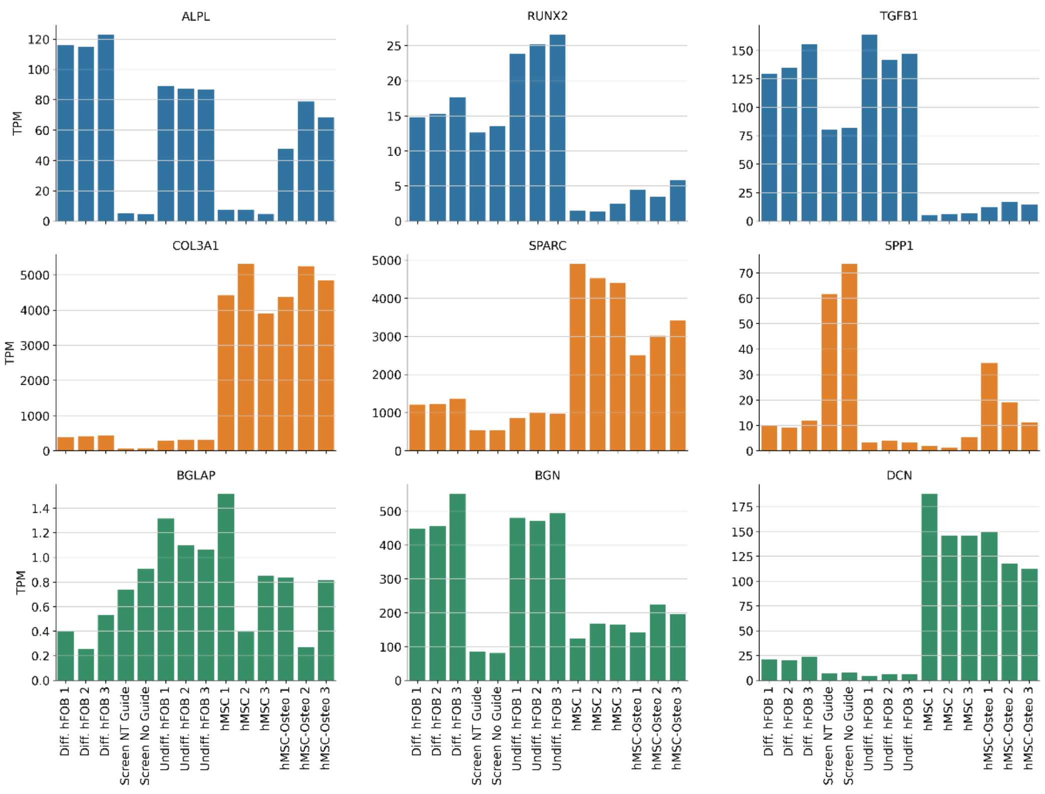


**Supplementary Fig. 6 - Comparison of Untargeted Screen Cells and Bulk Cell Models.**

(A) Principal components analysis of bulk RNA-seq of differentiated and undifferentiated hFOBs, hMSCs, and hMSC-Osteoblasts and pseudo-bulk scRNA-seq of dCAS9-KRAB-transfected hFOBs collected from the CRISPRi screen. Screen cells either received no guide or only non-targeting (NT) guides. (B) Osteoblast proliferation (blue), maturation (orange), and mineralization (green) marker gene expression in the same cells.

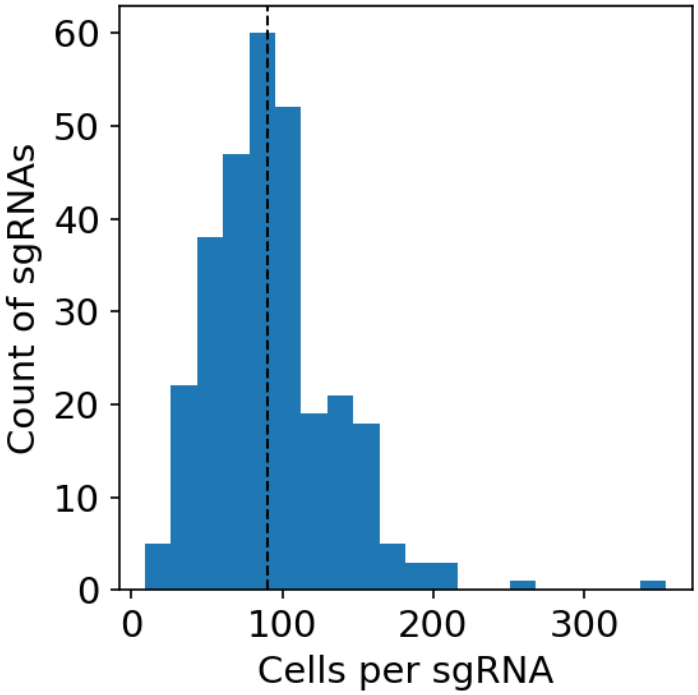


**Supplementary Fig. 7** - **Histogram of Cells per sgRNA given 27,385 Cells Retained after QC Filters**.

Median of 90 cells per sgRNA is labeled by the dashed line. The minimum number of cells per sgRNA is 9; and the maximum, 354.

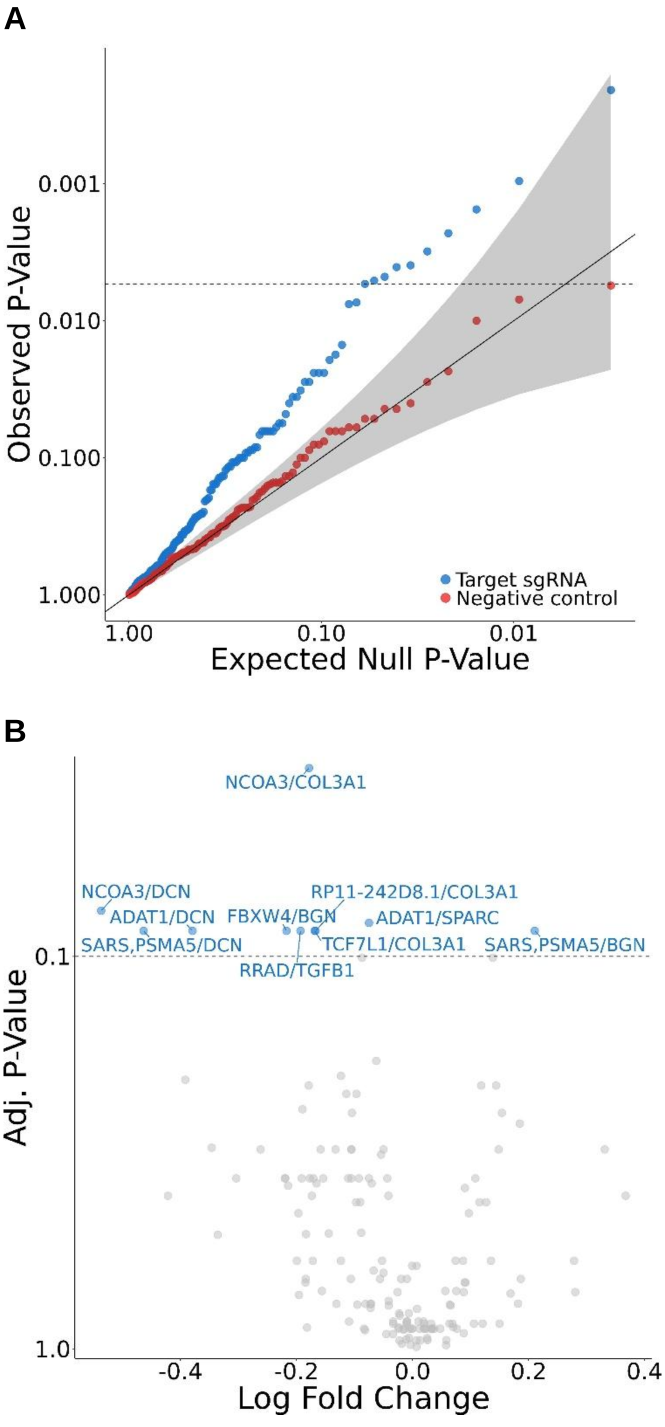


**Supplementary Fig. 8 - Trans-Perturbations of Osteoblast Marker Genes from hFOB CRISPRi Screen.**

(A) Quantile-quantile plot of SCEPTRE trans-perturbation analysis testing effect of 20 successful cis-perturbations on osteoblast marker genes. (B) Volcano plot of osteoblast-marker gene trans-perturbations. Each significant trans-perturbation is labeled in blue first by the gene(s) found to be perturbed in cis from the sgRNA and then, following the slash, by the perturbed osteoblast marker gene.

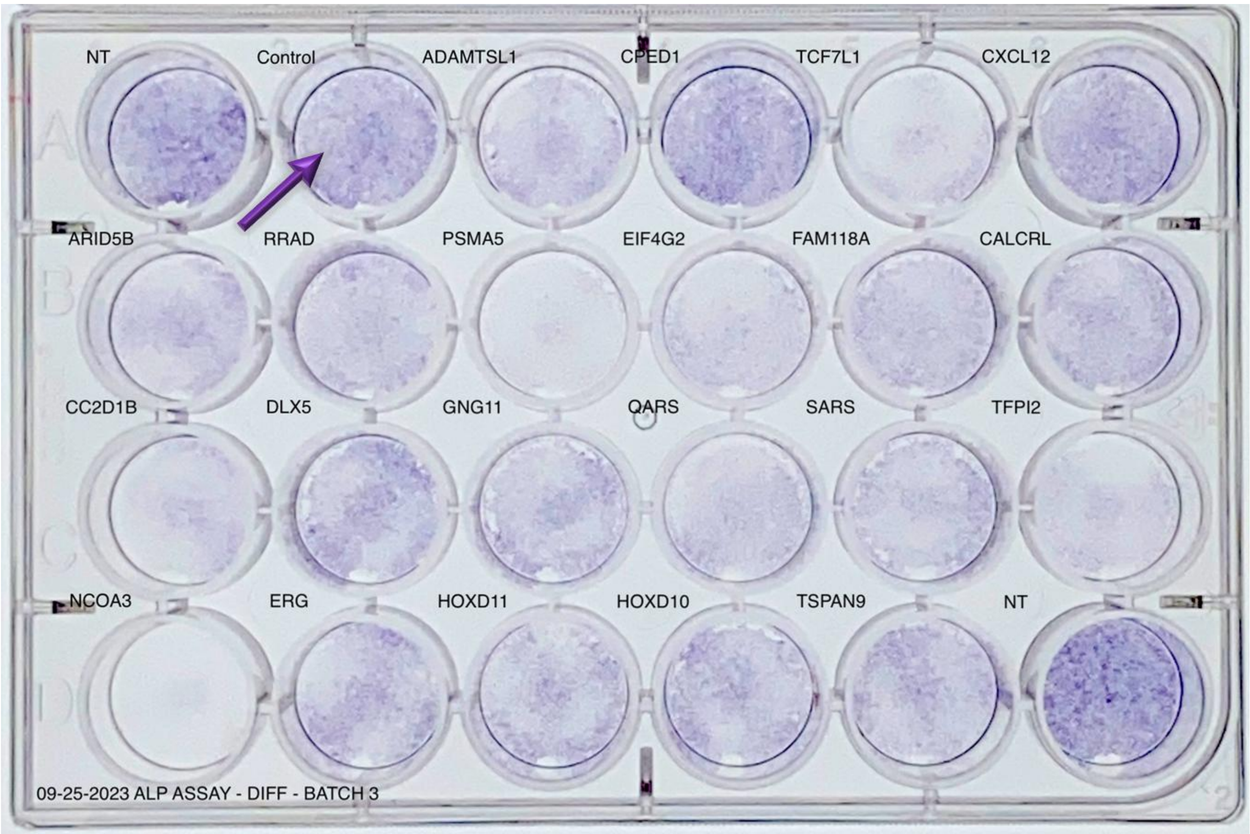


**Supplementary Fig. 9 - Representative hFOB ALP Assay Results.**

The purple arrow denotes scrambled control siRNA. Darker purple coloring of wells indicates higher levels of ALP secretion. NT wells contained no siRNAs.

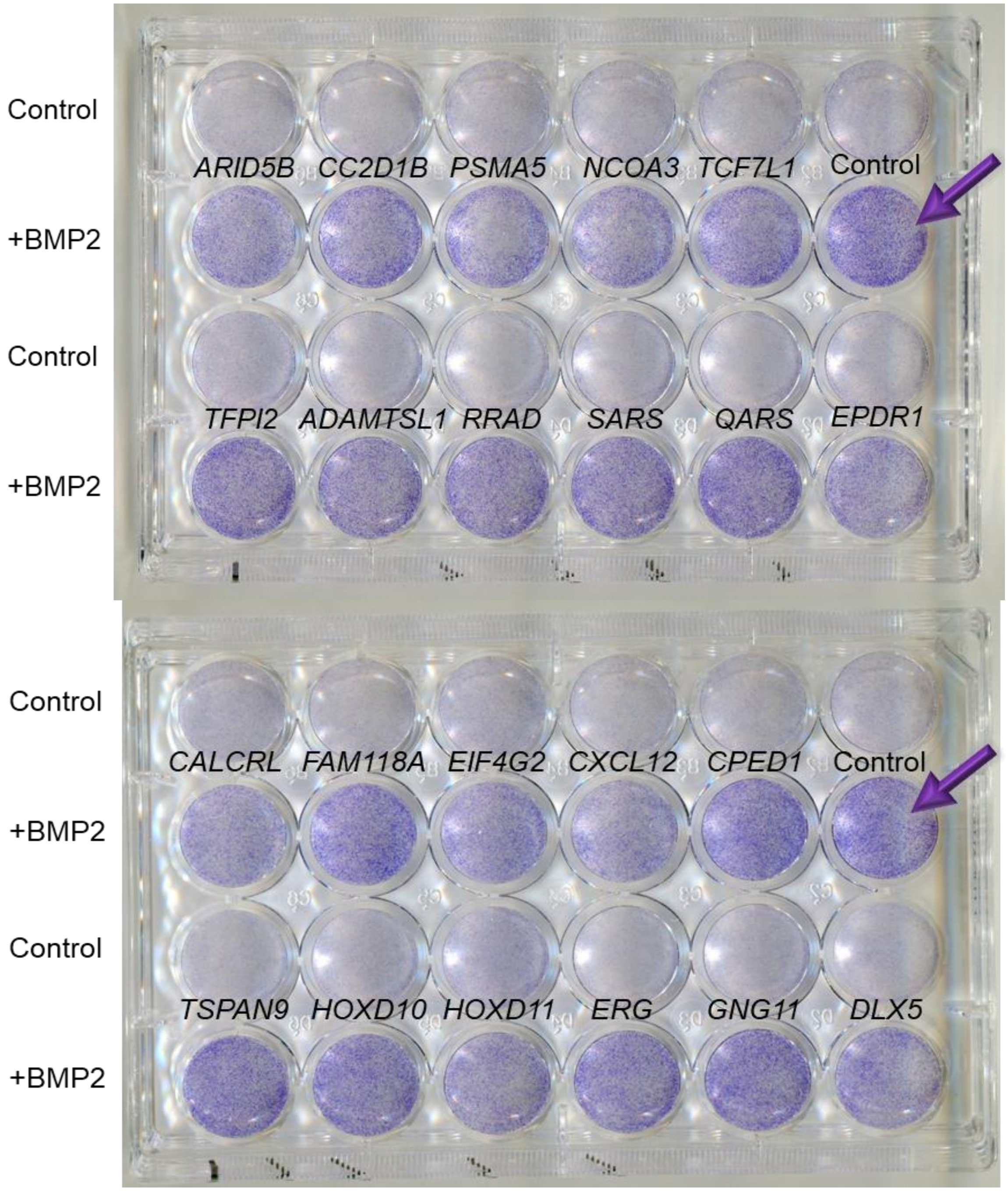


**Supplementary Fig. 10 - Representative hMSC-Osteoblast ALP Assay Results.**

Wells treated with scrambled control siRNAs and BMP2-supplemented media were used for plate normalization and are denoted by a purple arrow. Darker purple coloring of wells indicates higher levels of ALP secretion. *EPDR1* wells are not relevant to this work and are included only to present unmodified images.

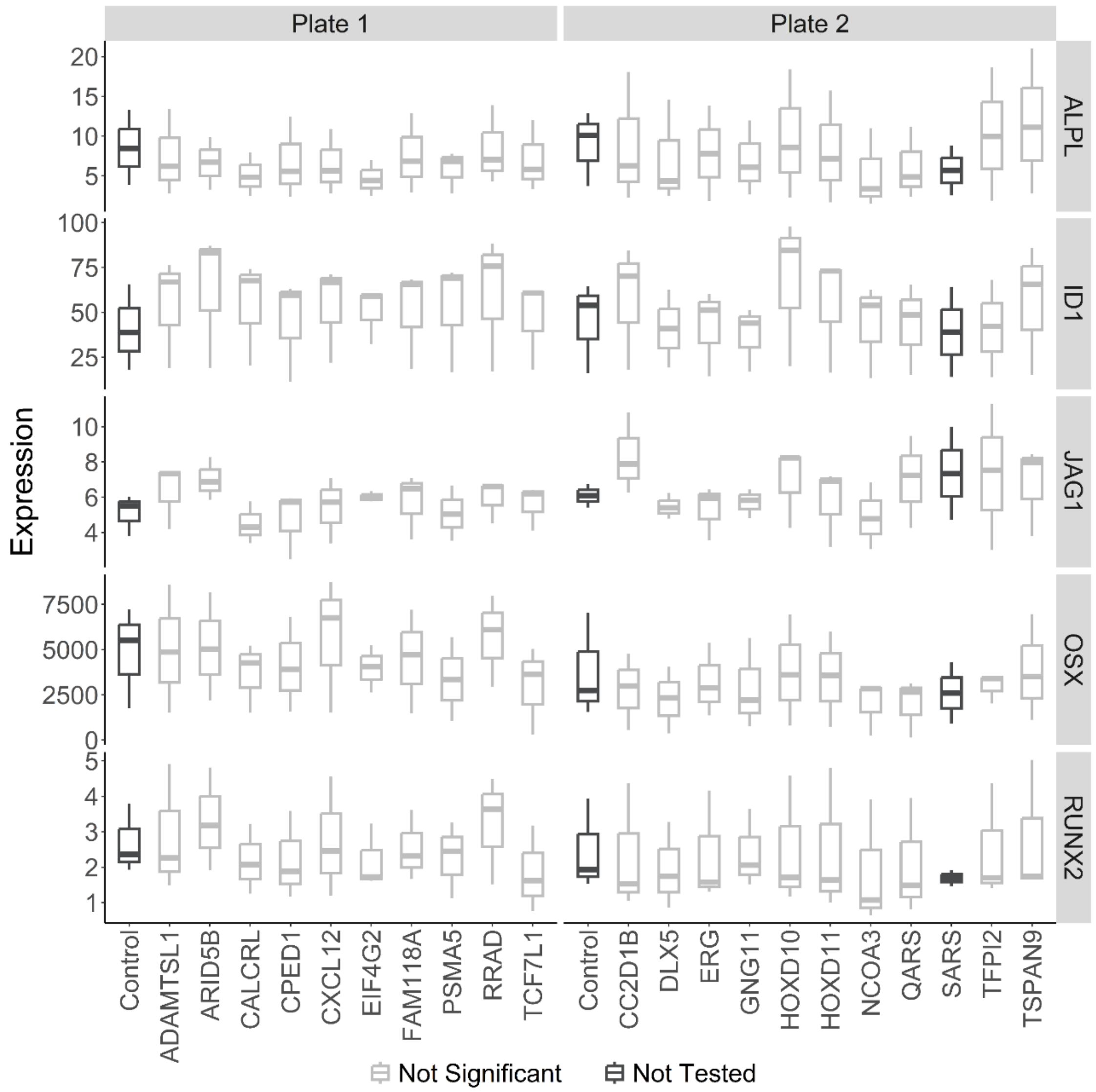


**Supplementary Fig. 11 - qPCR of Osteoblast Marker Gene Expression upon siRNA Knockdown.**

The boxplots present qPCR-measured expression of osteoblast marker genes in differentiated hMSC-osteoblasts under different siRNA knockdown conditions. Significant gene knockdowns (Benjamini-Hochberg adjusted *P*-value < 0.05) are plotted in blue and all tests are calculated relative to matched plate control siRNAs.

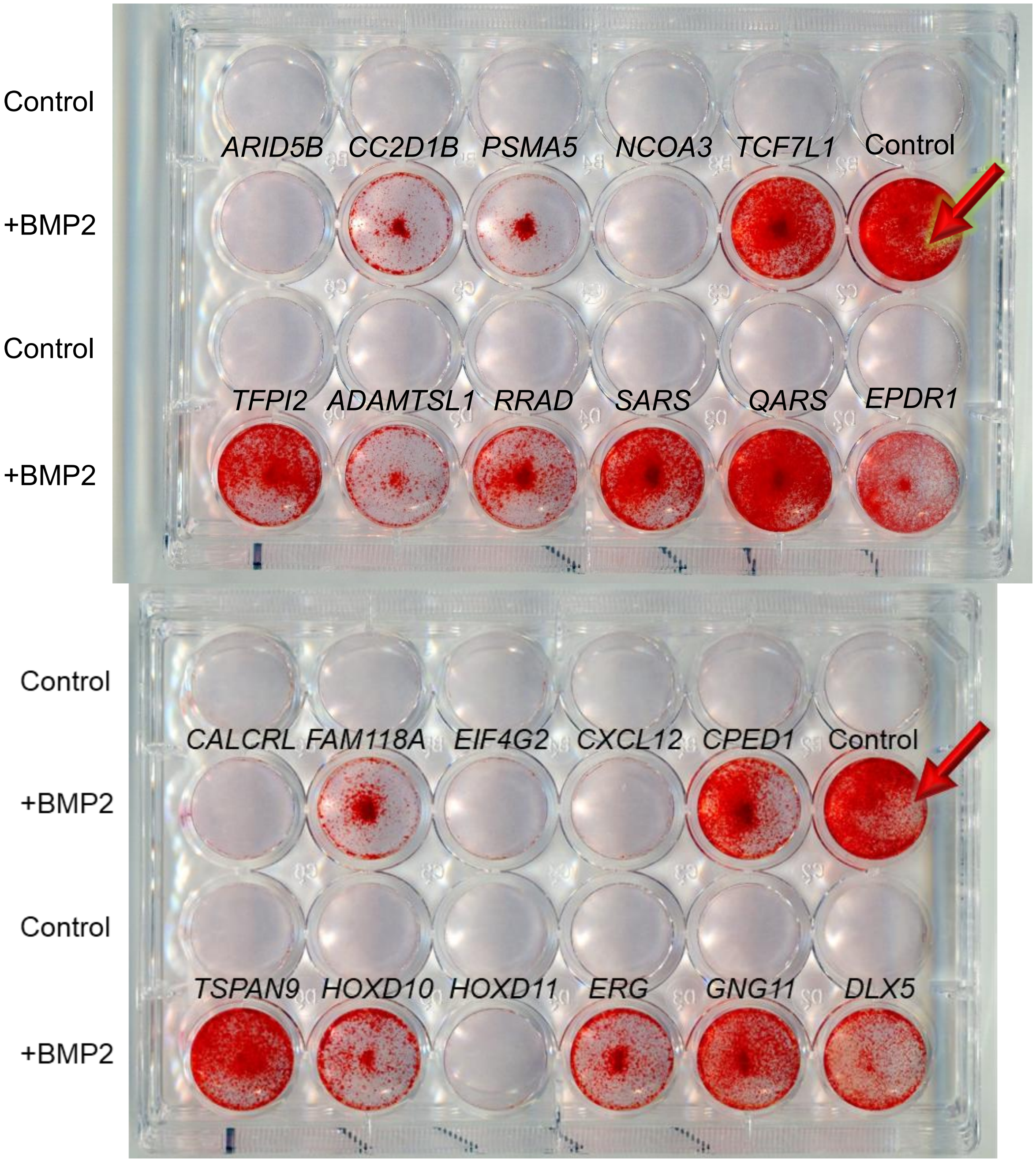


**Supplementary Fig. 12 - Representative hMSC-Osteoblast ARS Assay Results.** Wells treated with scrambled control siRNAs and BMP2-supplemented media were used for plate normalization and are denoted by a red arrow. Darker red coloring of wells indicates higher levels of mineral secretion. *EPDR1* wells are not relevant to this work and are included only to present unmodified images.

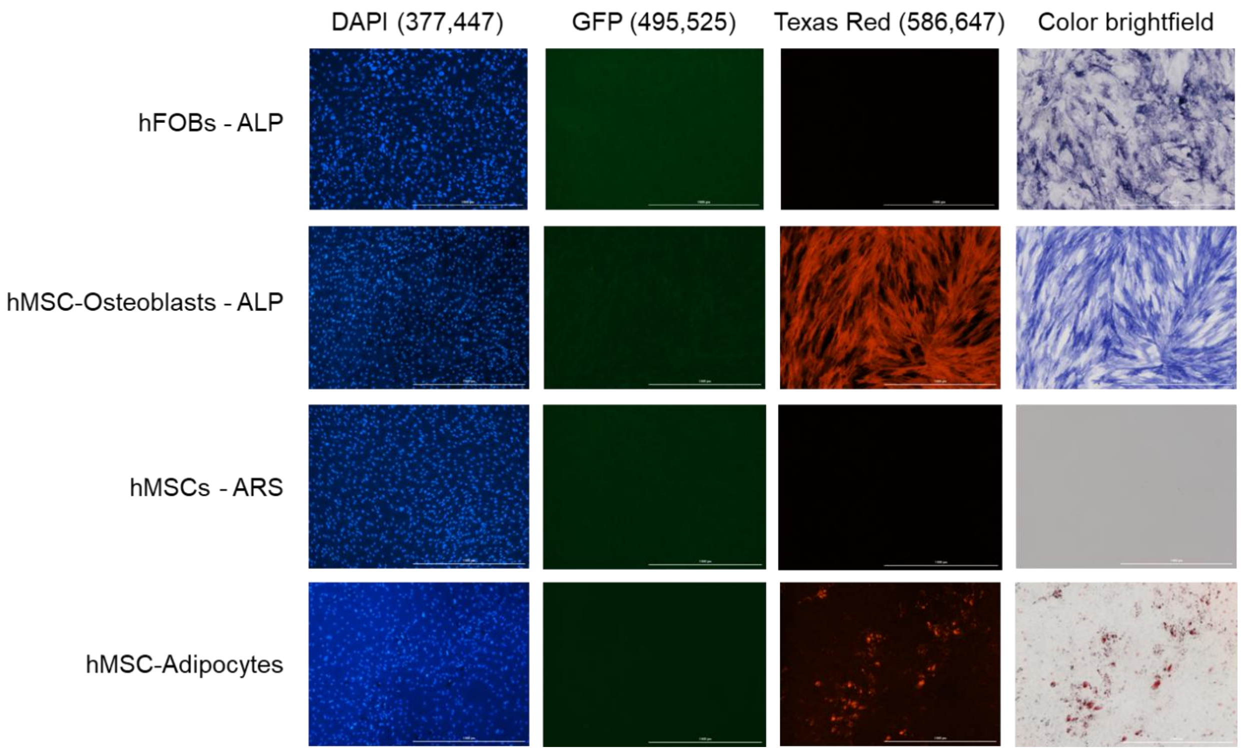


**Supplementary Fig. 13 - Representative DAPI Staining Images.**

Representative microscopy images shown for four functional assays on three fluorescence channels and brightfield. Cells were fixed in differentiated states prior to staining and imaging. All images shown were for scrambled, control siRNAs.

A

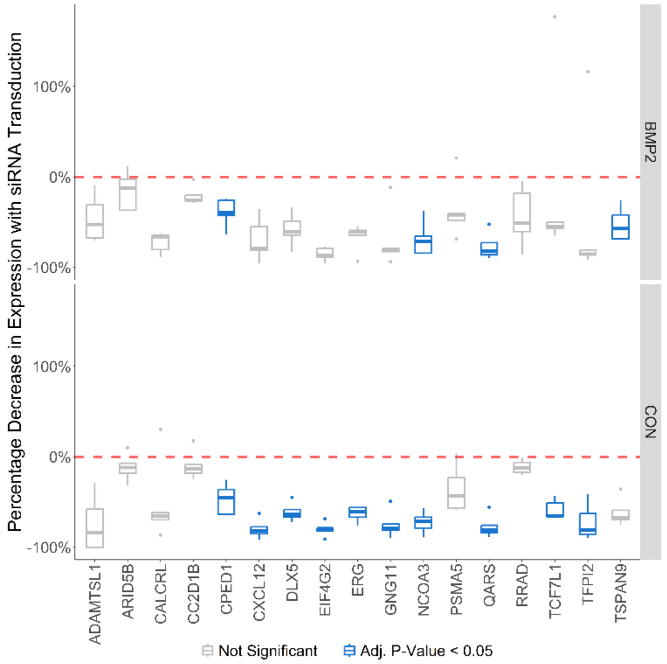


B

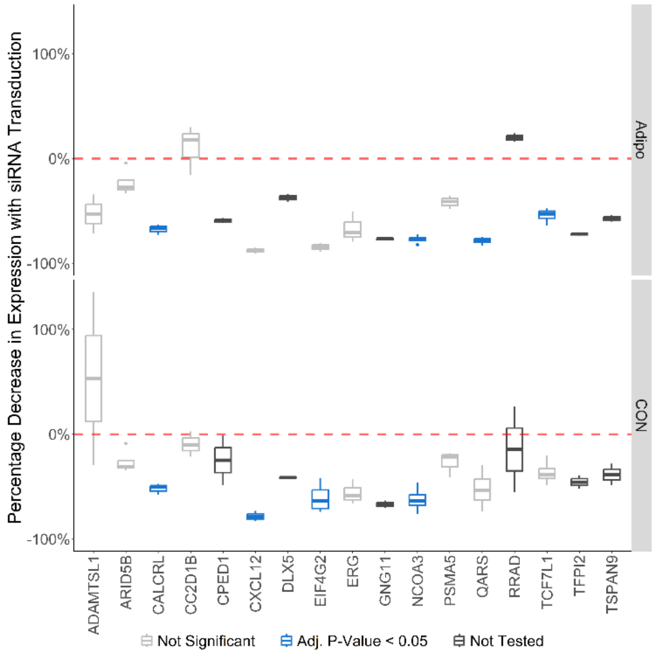


**Supplementary Fig. 14 - qPCR of Candidate Gene Expression upon siRNA Knockdown.**

The boxplots present percent changes between scrambled siRNA controls and gene-targeting siRNAs for expression of the targeted genes. Results are plotted separately by treatment status for (A) induced osteoblast differentiation and (B) adipocyte differentiation. Significant gene knockdowns (Benjamini-Hochberg adjusted P-value <

0.05) are plotted in blue and untested knockdowns with fewer than three biological replicates are plotted in charcoal.

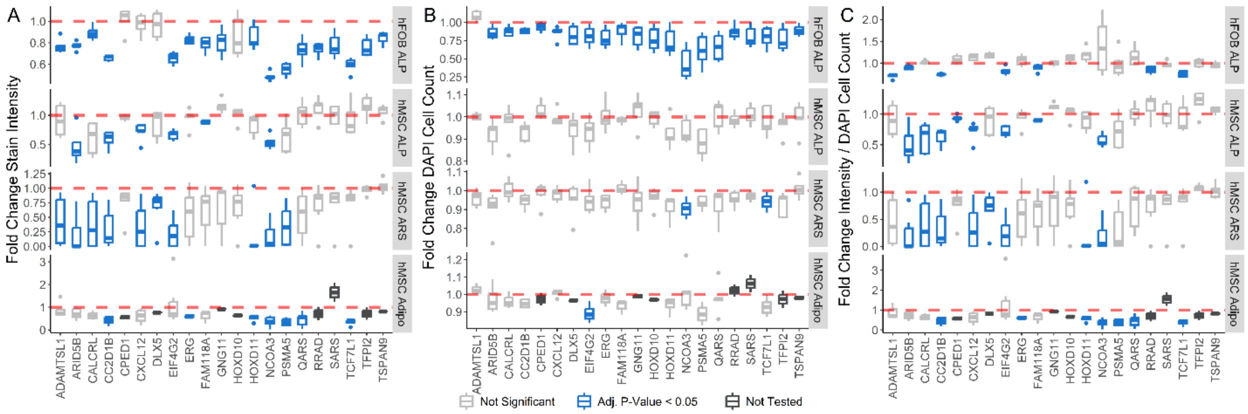


**Supplementary Fig. 15 - Plate-normalized assay results.**

Alkaline phosphatase (ALP) assay in hFOBs (top panels), ALP assay in hMSC-Osteoblasts (second from top), alizarin Red S (ARS) assay in hMSC-Osteoblasts (third from top), and adipogenesis assay in hMSCs (bottom) separated by (A) stain intensity, (B) cell counts, and (C) per-cell stain intensity. In each plot siRNA targets are listed along the x-axis and fold changes in the corresponding measurements along the y-axis. hMSC-based assays were measured and normalized in the treated state (with BMP2 or adipogenic induction media). Any siRNAs resulting in significant decreases of assayed measurements are displayed in blue and siRNAs with < 3 replicates were untested and are marked in charcoal.

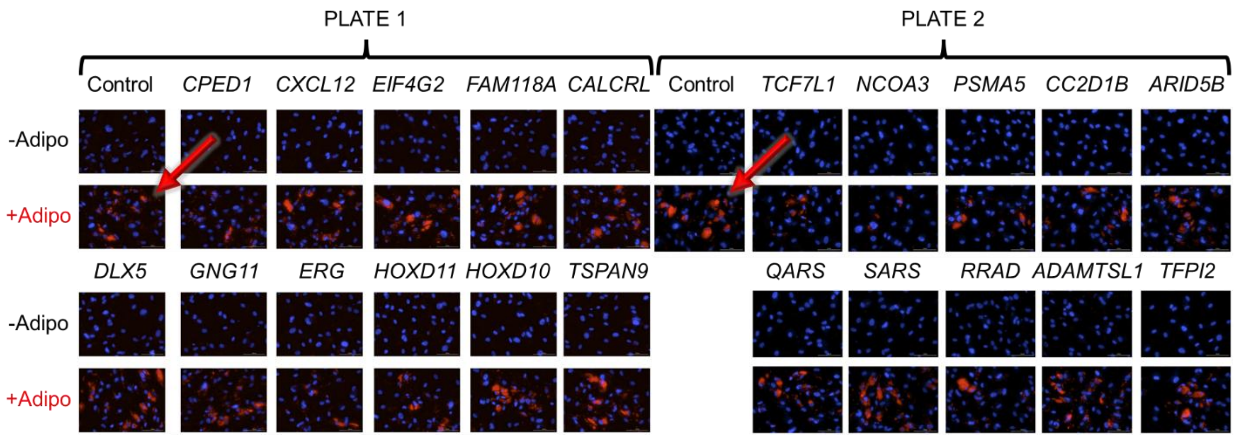


**Supplementary Fig. 16 - Representative hMSC Adipogenesis Assay Results.**

Images taken under 20x magnification with DAPI staining (blue) to indicate the position of nuclei and Oil Red O staining (red) to indicate lipid droplets. Cells treated with scrambled control siRNAs and adipogenic media were used for plate normalization and are denoted by a red arrow.

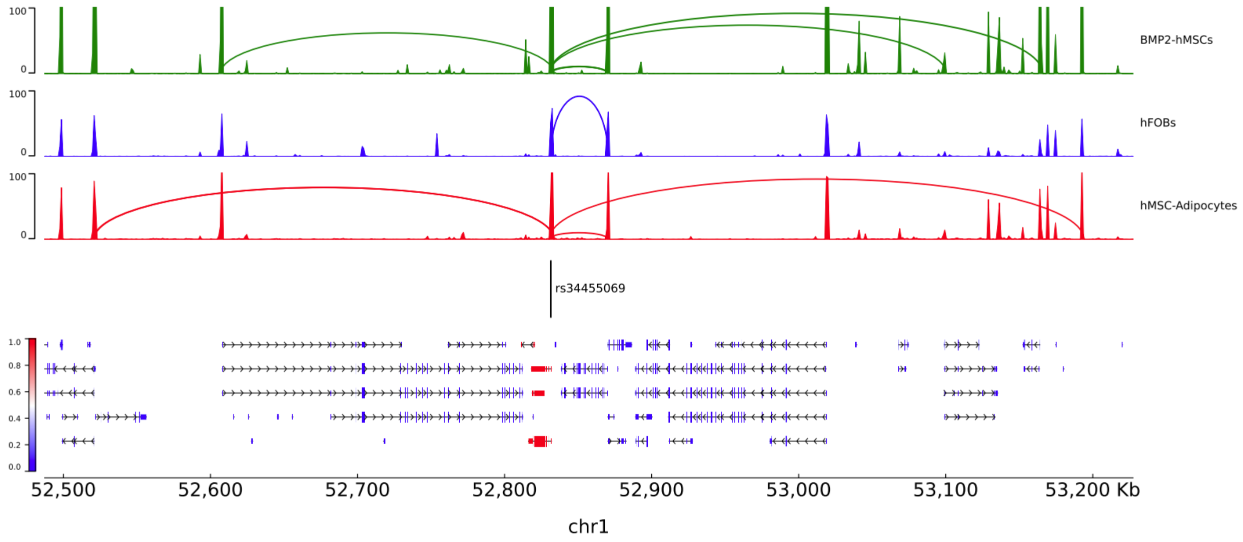


**Supplementary Fig. 17 - pyGenomeTracks plots of the *CC2D1B* locus.**

ATAC-seq read and Capture-C chromatin loops measured in hMSC-Osteoblasts, hFOBs, and hMSC-adipocytes are plotted above. Targeted SNPs and genes are plotted below. All isoforms of the perturbed gene in the CRISPRi screen, *CC2D1B*, are colored red.

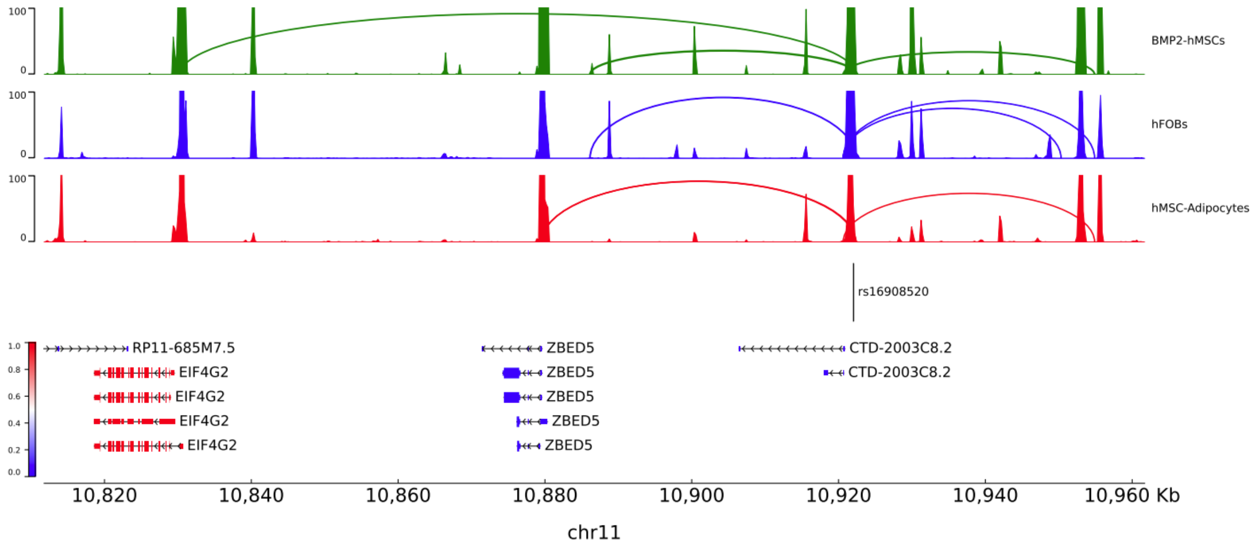


**Supplementary Fig. 18 - pyGenomeTracks plots of the *ZBED5-AS1* locus.**

ATAC-seq read and Capture-C chromatin loops measured in hMSC-Osteoblasts, hFOBs, and hMSC-adipocytes are plotted above. Targeted SNPs and genes are plotted below. All isoforms of genes perturbed in the CRISPRi screen are colored red.

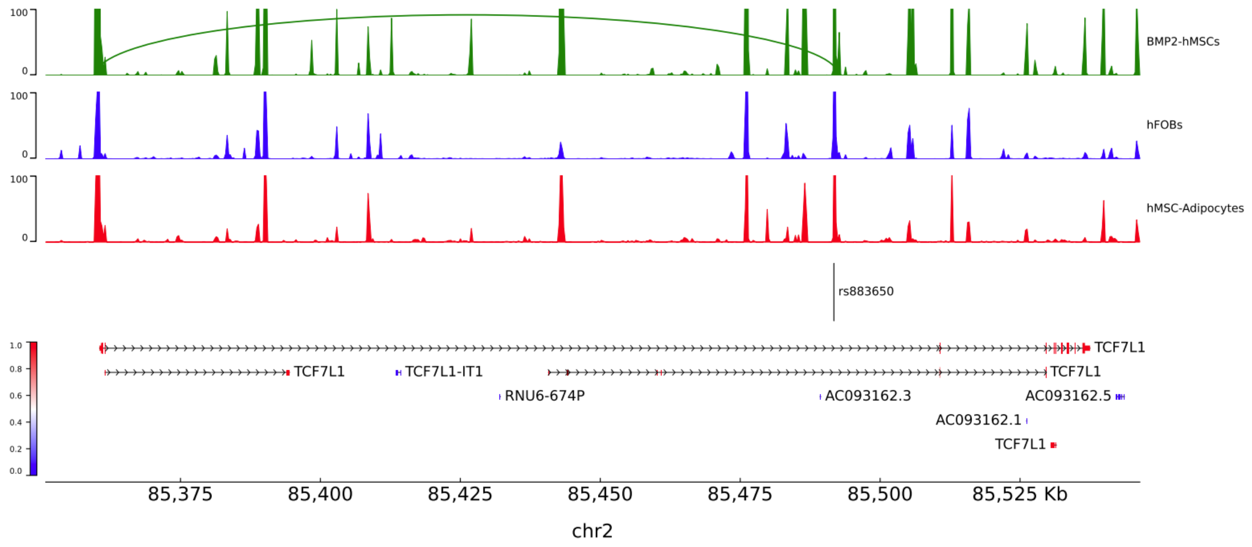


**Supplementary Fig. 19 - pyGenomeTracks plots of the *TCF7L1* locus.**

ATAC-seq read and Capture-C chromatin loops measured in hMSC-Osteoblasts, hFOBs, and hMSC-adipocytes are plotted above. Targeted SNPs and genes are plotted below. All isoforms of genes perturbed in the CRISPRi screen are colored red.

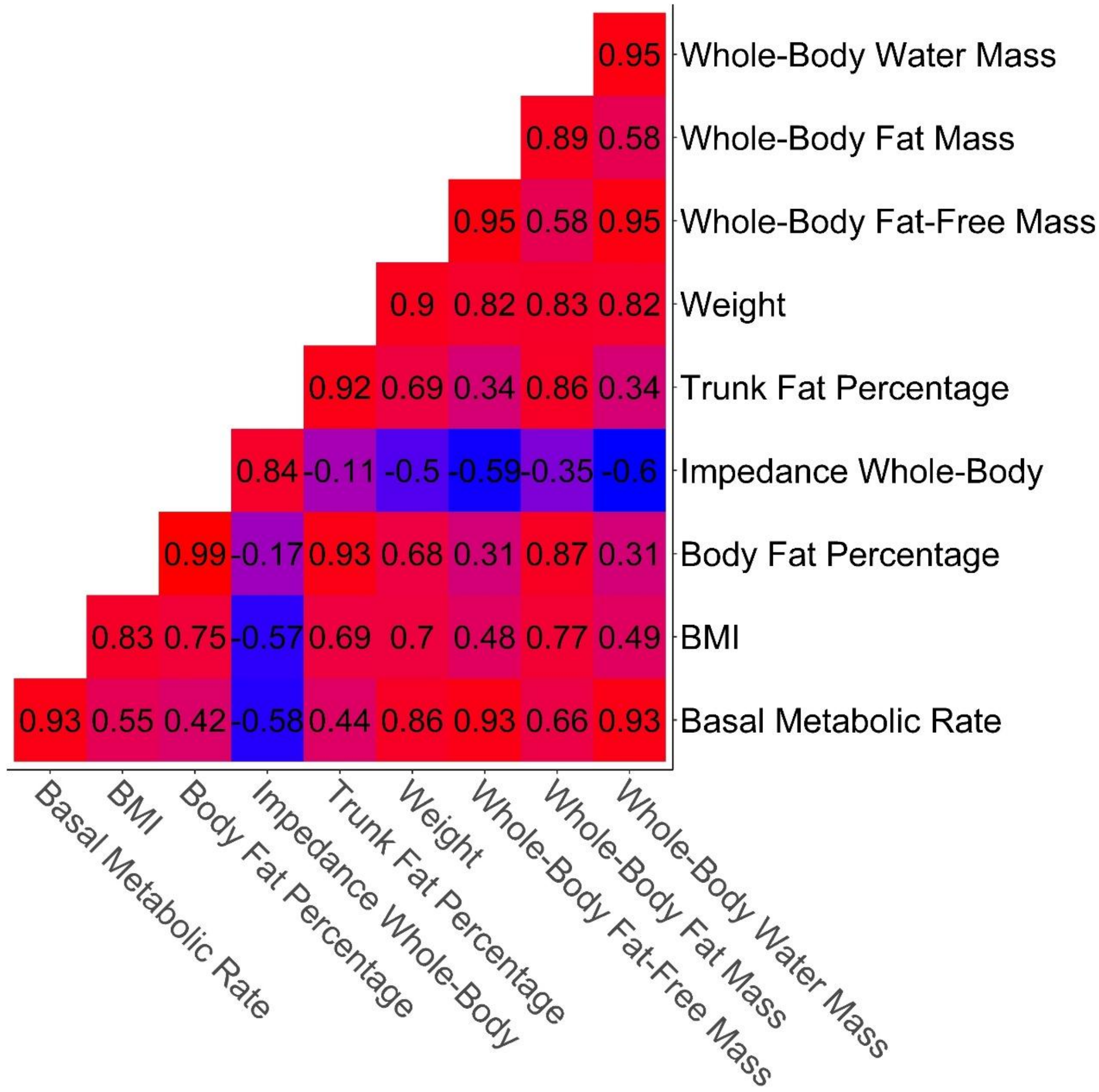


**Supplementary Fig. 20 - Significant Genetic Correlations between Impedance and Weight Derived Traits.**

Heatmap of genetic correlations between nine weight and impedance-based traits.

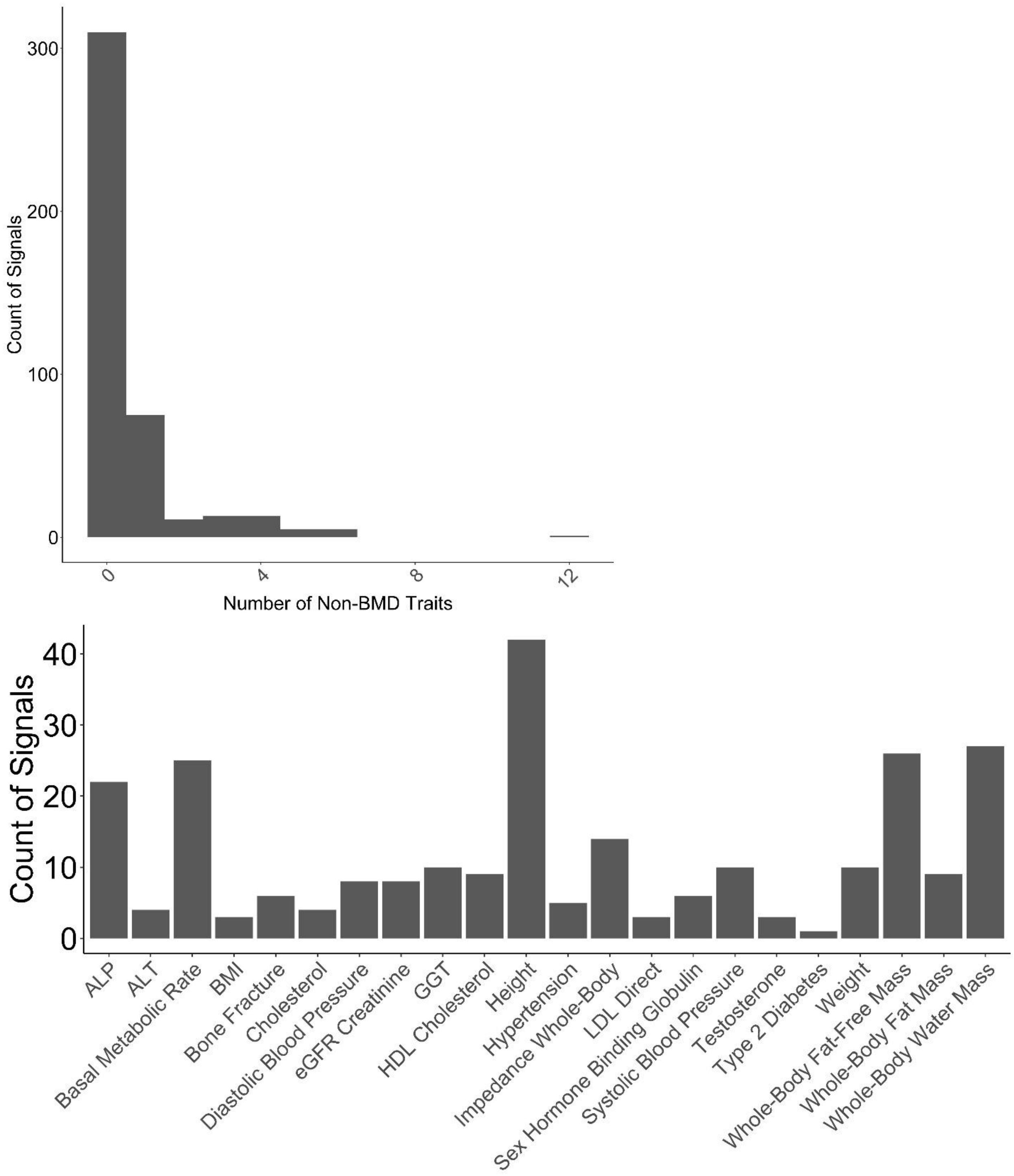


**Supplementary Fig. 21 - Distributions of Fine-Mapped BMD Signals and Traits.** The top histogram reflects the number of non-BMD traits mapped to each of the 433 identified BMD signals. The bottom bar plot shows counts of the number of signals mapped to each of 22 traits with one or more signals.

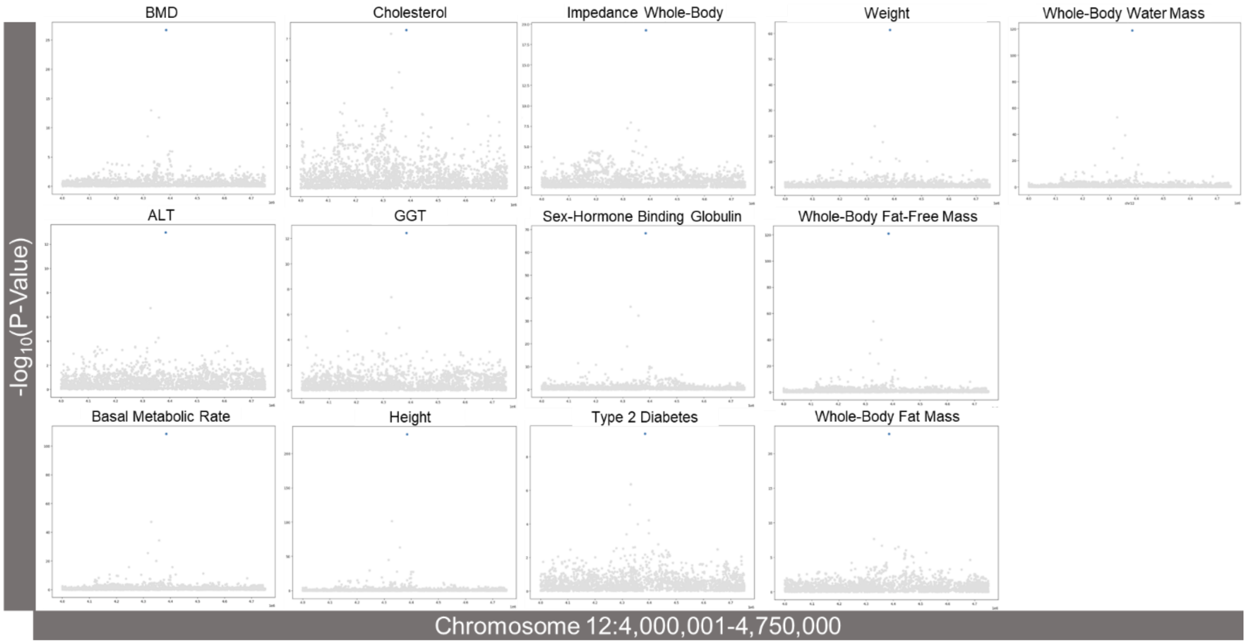


**Supplementary Fig. 22 - CAFEH Fine-Mapping Results of *CCND2* Locus.**
One credible set containing rs76895963 was identified and is marked in blue.

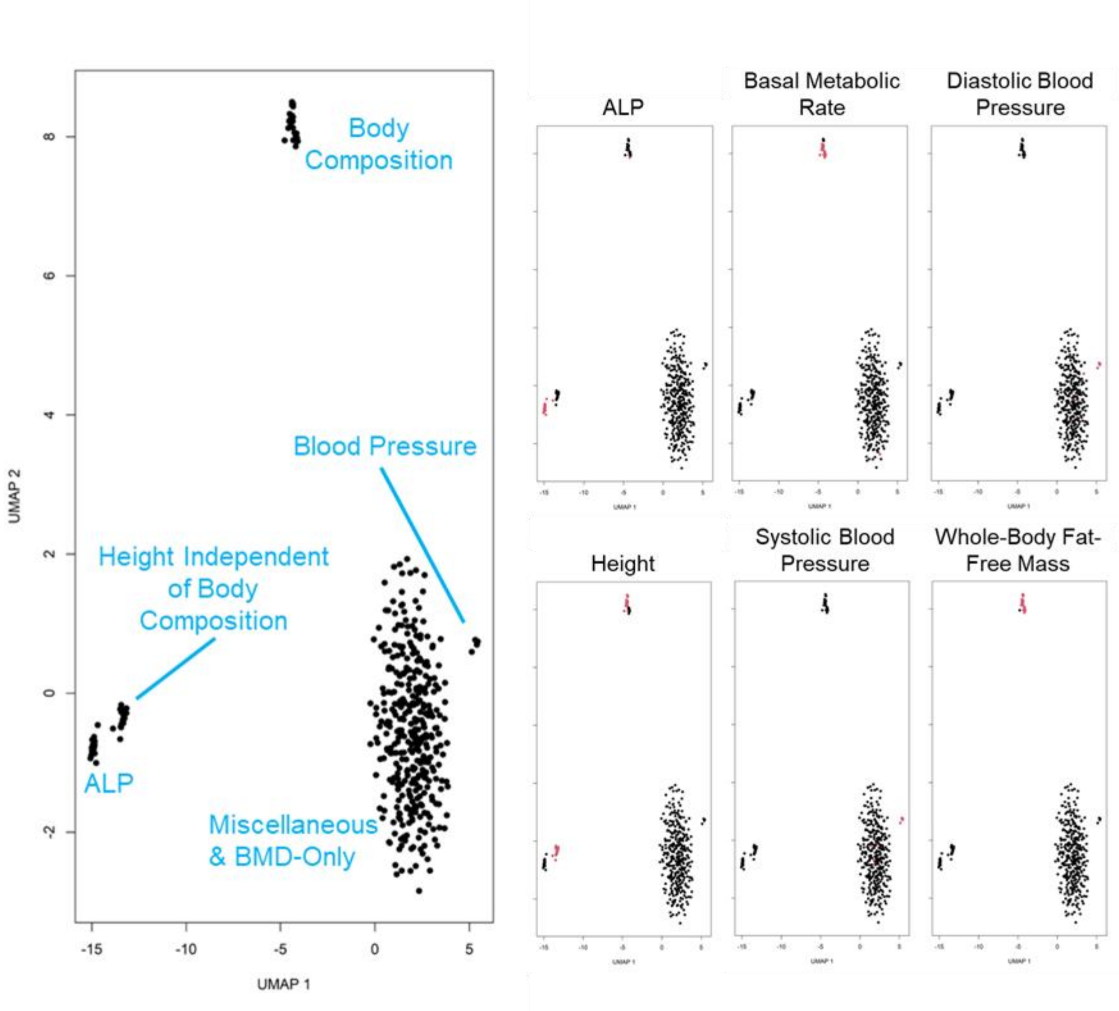


**Supplementary Fig. 23 - UMAP Plots of 433 BMD Signals.**

The right six plots are colored by signal sharing with red points representing signals that are considered shared by the selected trait. The left most plot has been manually labeled to reflect the predominant traits in each of the observed clusters.

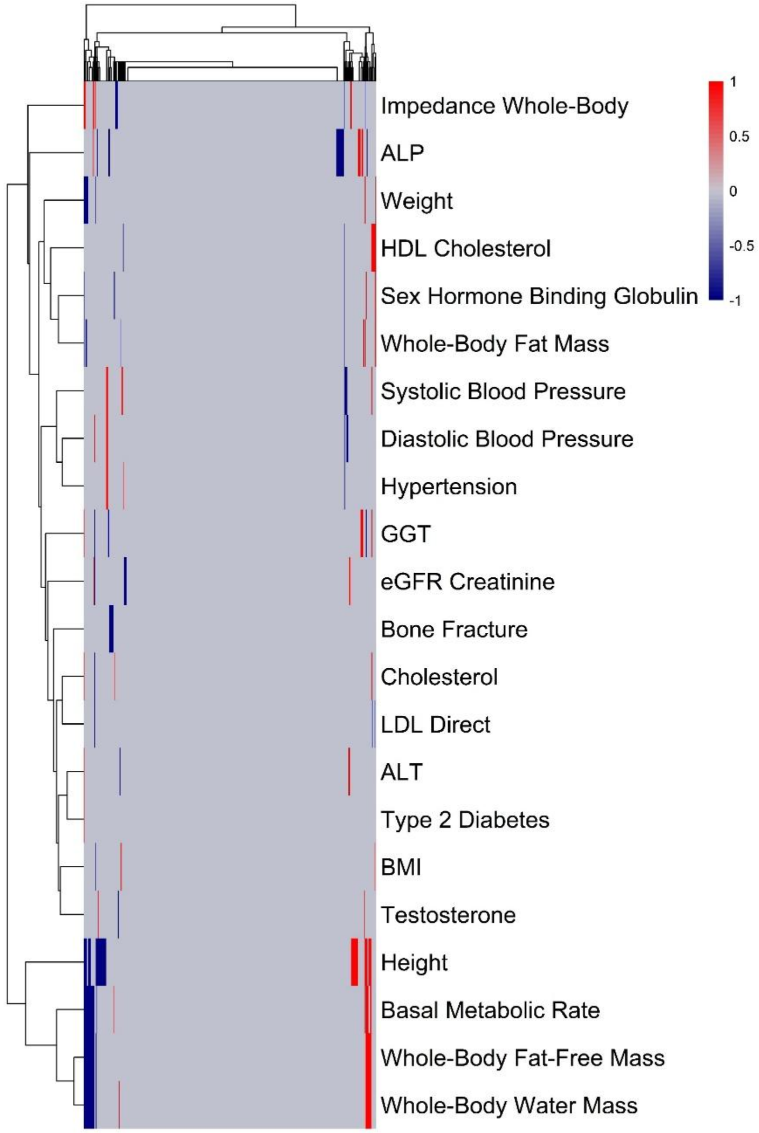


**Supplementary Fig. 24 - Heatmap and Dendrogram of 433 BMD Signals Split by Effect Direction.**

Traits are plotted down the y-axis with individual signals plotted along the x-axis. Red indicates when a signal modulates the corresponding trait and the effect direction positively correlates with the effect on BMD, and blue indicates negative correlation. Signals and traits are clustered hierarchically using the complete-linkage method with Euclidean distances.

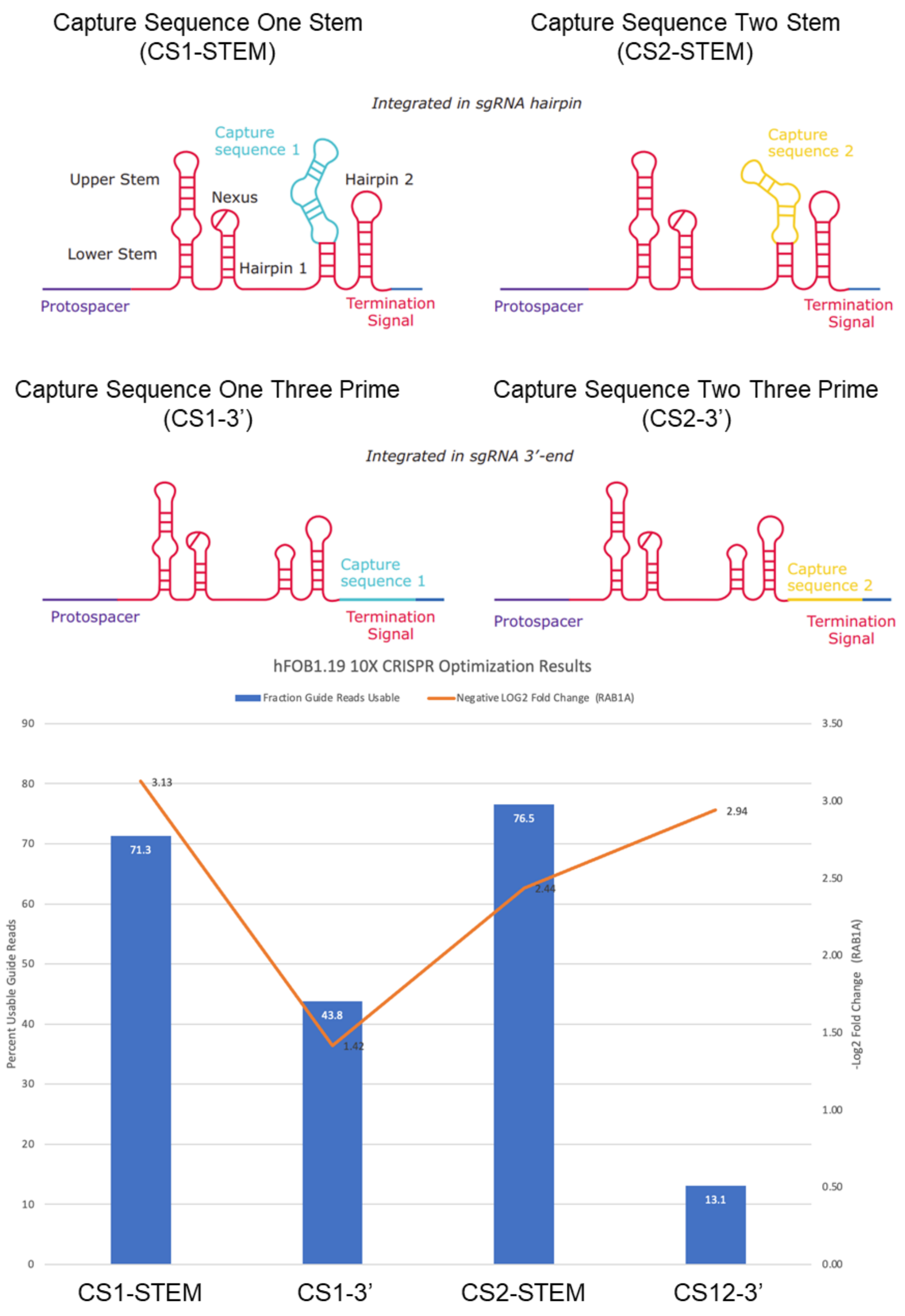


**Supplementary Fig. 25 - CRISPRi sgRNA Designs and hFOB Optimization Assay Results.**

Top panel shows possible Sigma-Aldrich sgRNA designs provided by the manufacturer and the bottom plot shows the results of the optimization assay measuring the capture rate of the sgRNA and the expression of *RAB1A* after knockdown.

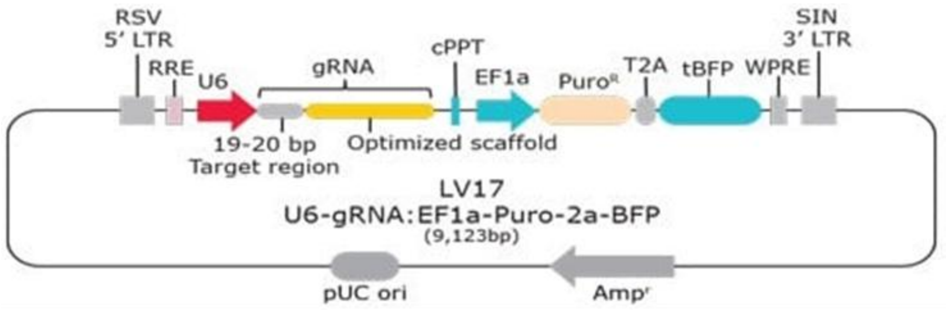


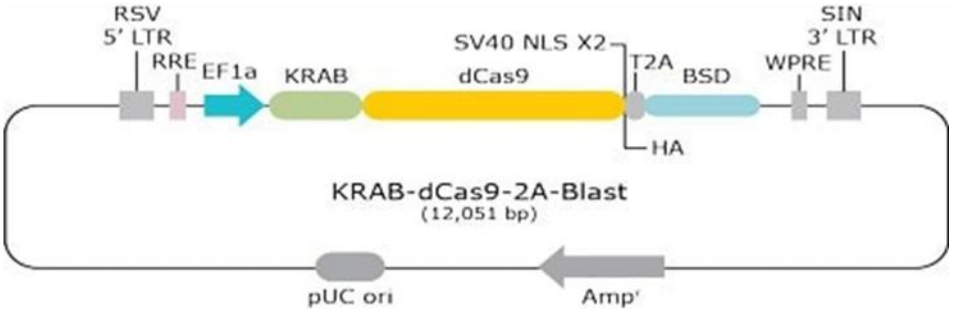


**Supplementary Fig. 26 - CRISPRi Lentiviral Construct Designs.**

Illustrations of Sigma-Aldrich sgRNA and KRAB-dCAS9 lentiviral vectors provided by the manufacturer.
